# A Long-Context Generative Foundation Model Deciphers RNA Design Principles

**DOI:** 10.64898/2026.03.17.712398

**Authors:** Yanjie Huang, Guangye Lv, Anyue Cheng, Wei Xie, Mengyan Chen, Xinyi Ma, Yijun Huang, Yueyang Tang, Qingya Shi, Jiahao Wang, Zining Wang, Junxi Wang, Yunpeng Xia, Lu Zhao, Yifang Cai, Jack X. Chen, Shuangjia Zheng

**Author notes:** These authors contributed equally to this work (Yanjie Huang, Guangye Lv).

## Abstract

Programmable design of RNA sequences with defined functions remains a central challenge in biology. Despite recent advances, existing RNA generative models lack robust controllable design capabilities and are constrained by short context windows, limiting their capacity to model the complex evolutionary manifold of full-length transcripts. Here, we introduce EVA (Evolutionary Versatile Architect), a long-context generative RNA foundation model trained on over 114 million full-length RNA sequences spanning the full breadth of evolutionary diversity. By integrating a Mixture-of-Experts architecture with an 8,192-token context window, EVA learns a unified, evolution-consistent representation of RNA sequence space that captures both global transcript structure and fine-grained functional elements. EVA establishes a unified framework for a broad spectrum of RNA design tasks within a single architecture, ranging from mutational fitness prediction to context-aware sequence engineering. We demonstrate EVA’s versatility through the de novo design and targeted optimization of diverse RNA classes, including tRNAs, aptamers, and CRISPR guide RNAs, as well as mRNA and circular RNA therapeutics. In systematic evaluations, EVA achieves state-of-the-art performance across seven of nine established benchmarks, improving structural modeling accuracy by up to an order of magnitude compared to existing approaches. EVA is fully open-source at the following repository: https://github.com/GENTEL-lab/EVA.

## 1 Introduction

RNA is a key molecular entity that performs essential functions within cells. It carries genetic instructions (mRNA) ^1^, folds into complex structures to carry out cellular roles (tRNA, rRNA) ^2^, catalyzes reactions (ribozymes) ^2^, and regulates gene expression (miRNA, lncRNA, snoRNA)^3,4^. This functional diversity positions RNA at the core of life process, driving evolutionary adaptation and cellular complexity ^5^. Consequently, the programmable design of RNA has become a central frontier for both decoding biological logic and engineering next-generation medicines.

The recent expansion of large-scale RNA sequence resources has therefore motivated RNA foundation models that aim to capture these evolution-imposed regularities from data and transfer them across tasks^6–12^. Much of the field has so far emphasized encoder-only, MLM-trained architectures that are particularly effective for representation learning and inference, whereas design-oriented goals are more naturally served by decoder-only generative modeling that can synthesize sequences with explicit control over context and constraints (Supplementary Table S1). Although early decoder-based RNA generators have begun to explore this direction, they typically remain limited in regime, either focusing on selected functional RNA classes ^13–15^ or operating with context lengths that are short relative to full transcripts ^16,17^, and the effective training signal can be further diluted by corpus heterogeneity, including datasets that interleave RNA and DNA. Moreover, in contrast to genome foundation models that have demonstrated scalability in the DNA domain ^18,19^, RNA modeling has yet to exhibit comparable scaling behavior, likely due to weaker sequence conservation and the fragmented nature of available RNA datasets. These challenges motivate the development of scalable, long-context generative RNA foundation models capable of capturing the complex evolutionary manifold of full-length transcripts to enable controllable design across diverse RNA classes and taxa.

Herein, we present EVA (Evolutionary Versatile Architect), a long-context generative RNA foundation model trained on 114 million full-length RNA sequences (231.3B nucleotides) spanning structural, regulatory, coding, and viral RNAs across the tree of life. Built on a 1.4B-parameter decoder-only Transformer with a Mixture-of-Experts backbone and an 8,192-token context window, EVA jointly optimizes autoregressive generation and masked infilling, enabling both *de novo* sequence synthesis and context-constrained redesign. Explicit conditioning on RNA type and taxonomic lineage allows it to disentangle universal RNA grammar from lineage-specific adaptations.

Across mutation-effect prediction, conditional generation, and functional RNA design tasks, EVA consistently outperforms prior RNA foundation models, supporting controllable design across 11 major RNA classes without task-specific fine-tuning. It achieves state-of-the-art zero-shot fitness prediction for ncRNAs and mRNAs, and extends to DNA and protein in exploratory settings. Sampling from its evolution-consistent RNA manifold yields sequences that are structurally plausible yet evolutionarily novel, with a more than 10-fold improvement in structural modeling accuracy. To support the community, we release the full training corpus (OpenRNA v1), EVA model checkpoints at all scales, detailed training protocols, and an interactive web server. By integrating scalable generative modeling, high-performance predictive scoring, and interpretable latent representations within a single framework, EVA provides a foundation for systematic RNA exploration and design across evolutionary diversity.

## 2 Results

### 2.1 EVA model architecture, training procedures, data, and evaluations

EVA scales to 1.4 billion parameters to unify the previously disparate domains of non-coding RNA (ncRNA) and messenger RNA (mRNA) modeling. Equipped with an 8,192-token context window, EVA serves as a versatile engine that supports both Causal Language Modeling (CLM) and Generalized Language Modeling (GLM), enabling controllable generation conditioned on RNA family and taxonomic lineages (Figure 1A). To power this system, we introduce OpenRNA v1, a comprehensive RNA atlas totaling 114 million curated full-length sequences. Spanning diverse taxonomic groups, the dataset exhibits a broad distribution of sequences across the tree of life (Figure 1B), with phylum-level coverage across both prokaryotic and eukaryotic lineages (Figure S1).

**Figure 1:**
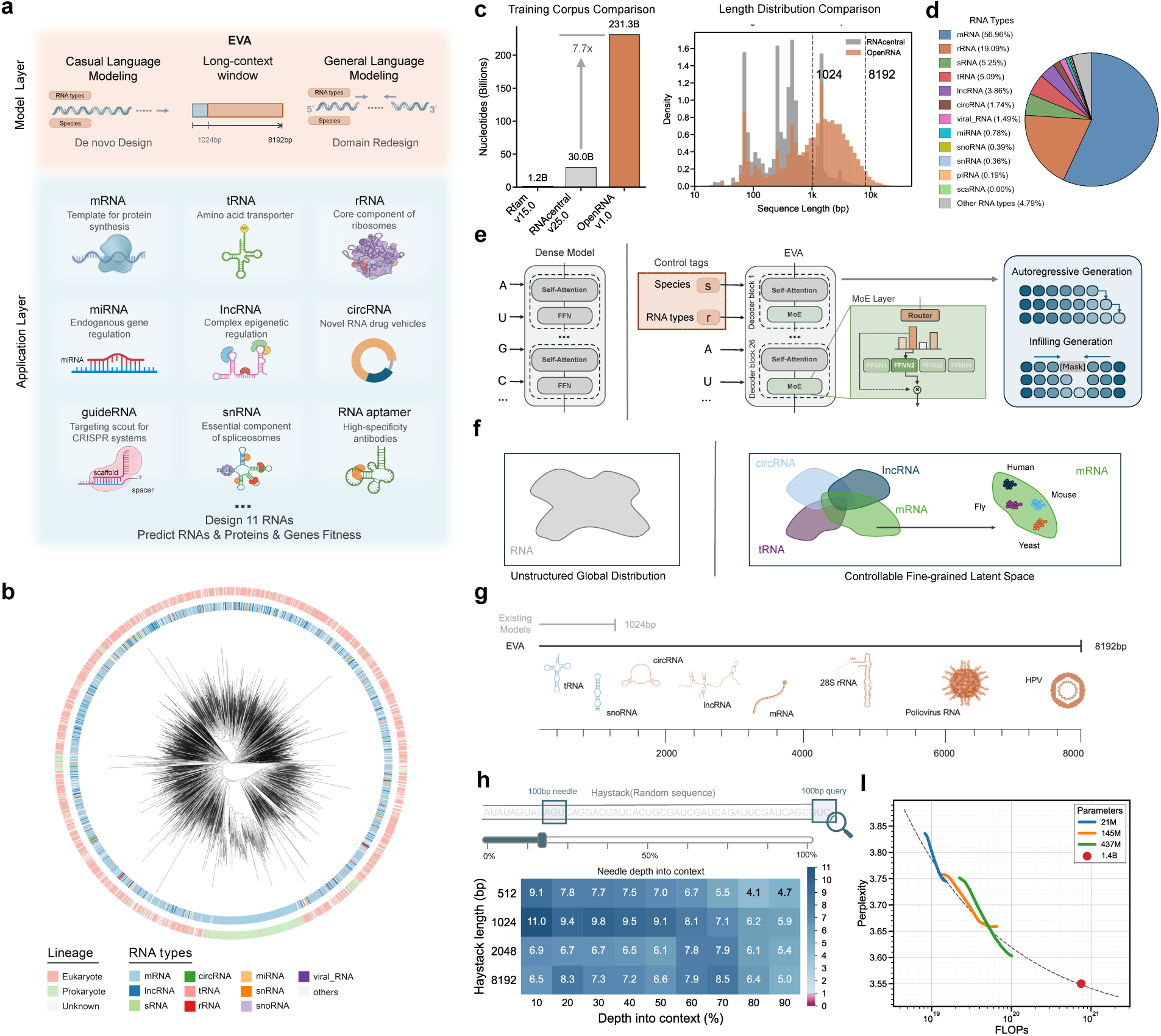
Overview of EVA data, model, and training. (a) Overview of the EVA model capabilities, and representative applications. (b) OpenRNA v1 comprises 114M curated full-length RNA sequences spanning all domains of life; Phylogenetic tree construction of training data. (c) Comparison of training data scale with prior work (left), and sequence-length distributions together with the number of full-length RNAs covered by different context windows (right). (d) RNA type distribution in OpenRNA v1. (e) EVA model architecture and key differences from prior RNA language models: Mixture-of-Experts (MoE) architecture, support for both CLM and GLM generation, and conditioning on RNA type and taxonomic labels. (f) Illustration of the unstructured global distribution learned by conventional RNA LMs versus the controllable fine-grained latent space learned by EVA. (g) Comparison of context window lengths between EVA (8,192 nt) and prior RNA language models (1,024 nt), highlighting broader RNA design coverage enabled by long context. (h) Long-context “needle-in-a-haystack” evaluation demonstrating EVA’s ability to recall long-range dependencies under an 8,192-token context window. (i) Scaling behavior showing decreasing perplexity with increasing model size, indicating that larger models better capture RNA sequence grammar.

To capture the hierarchical grammar of full-length RNAs, we adopted an 8,192-token context window, an 8-fold increase over conventional RNA models, expanding the designable RNA space (Figure 1C). This extended context enables the model to leverage our large-scale corpus effectively, capturing long-range dependencies across diverse architectures and covering over 95% of known full-length transcripts (Figure 1C). The dataset integrates 15 distinct RNA categories, providing a comprehensive representation of the RNA landscape (Figure 1D).

To model this diversity, EVA employs a decoder-only Transformer with a Mixture-of-Experts (MoE) backbone (Figure 1E). MoE architectures have demonstrated superior parameter efficiency and performance over dense Transformers in protein language models ^20,21^; EVA is the first RNA foundation model to adopt this design, and we validate its advantage empirically (Supplementary Table S5). However, training at this scale reveals that RNA sequences exhibit lower evolutionary conservation and a more severely imbalanced cluster size distribution than proteins (Figure S2C), causing standard training to overfit on abundant sequences while neglecting rare functional families (Figure S3A). We therefore implemented an evolutionary conservation-based sampling strategy (see Methods), which rebalances RNA-type abundances (Figure S2D) and improves model generalization (Figure S3A), with square-root sampling achieving the best balance between coverage and cluster down-weighting (Figure S2C, S3B).

To fully leverage this curated data while enabling controllable generation, we devised a two-stage training curriculum that maintains both CLM and GLM objectives throughout. In the pre-training stage, we condition the model solely on RNA-type tags, which we found to significantly enhance the quality of generated sequences across all criteria (GC content validity, sequence completeness, and low repetitiveness) by providing explicit family cues that help disentangle the unstructured global distribution (Figure S5). This allows the model to learn fundamental RNA syntax and structural motifs without being overwhelmed by fine-grained taxonomic details. In the subsequent mid-training stage, we introduce lineage (taxonomic) tags. This sequential integration of conditional information is critical; we found that introducing species-level constraints too early leads to overfitting, whereas our staged approach enables the model to first internalize universal RNA grammar before refining its latent space for species-specific adaptability (Figure S3C). This strategy enables EVA to unify de novo generation and targeted region redesign within a single framework, transitioning from a generic global distribution to a controllable, fine-grained latent space tailored to specific biological contexts (Figure 1F).

We observe consistent scaling behavior across all trained model sizes (~21M to 1.4B), where perplexity decreases consistently with capacity, indicating progressively better capture of RNA grammar (Figure 1I). Hyperparameter exploration further reveals that mid-training lineage conditioning further reduces perplexity by disentangling species-specific constraints from universal RNA grammar (Figure S4B). Finally, using a “needle-in-a-haystack” evaluation, we demonstrate that EVA reliably detects perturbations as subtle as 1% introduced at arbitrary positions within sequences up to 8,192 tokens, confirming robust long-range dependency recall across its entire context window (Figure 1H).

### 2.2 EVA predicts mutational effects across RNA, DNA, and protein

Language models have been shown to capture fitness landscapes by assigning higher likelihood to functional sequences and lower likelihood to deleterious variants^18,22,23^. Since EVA is trained on full-length RNA sequences, including coding RNAs that simultaneously encode both transcript-level and protein-level constraints, its manifold naturally reflects selection pressures operating across multiple molecular scales. We therefore evaluate EVA likelihood as a unified scoring function across these modalities.

We benchmark EVA on diverse deep mutational scanning (DMS) assays covering both structured ncRNAs and mRNAs. For each assay, we compute the Spearman rank correlation between per-variant EVA log-likelihood scores and experimentally measured fitness effects. Across structured ncRNA families, including ribozymes, aptamers, and tRNAs, EVA achieves a mean Spearman correlation of 0.40, establishing state-of-the-art performance (Figure 2B). For mRNA assays, EVA attains a mean Spearman correlation of 0.31, surpassing prior RNA language models as well as dedicated codon optimization models, demonstrating sensitivity to translation efficiency and RNA stability constraints (Figure 2C). In both cases, EVA outperforms prior RNA/Codon language models trained on smaller or less phylogenetically diverse datasets, suggesting that scale and taxonomic breadth contribute to learning generalizable fitness landscapes.

**Figure 2:**
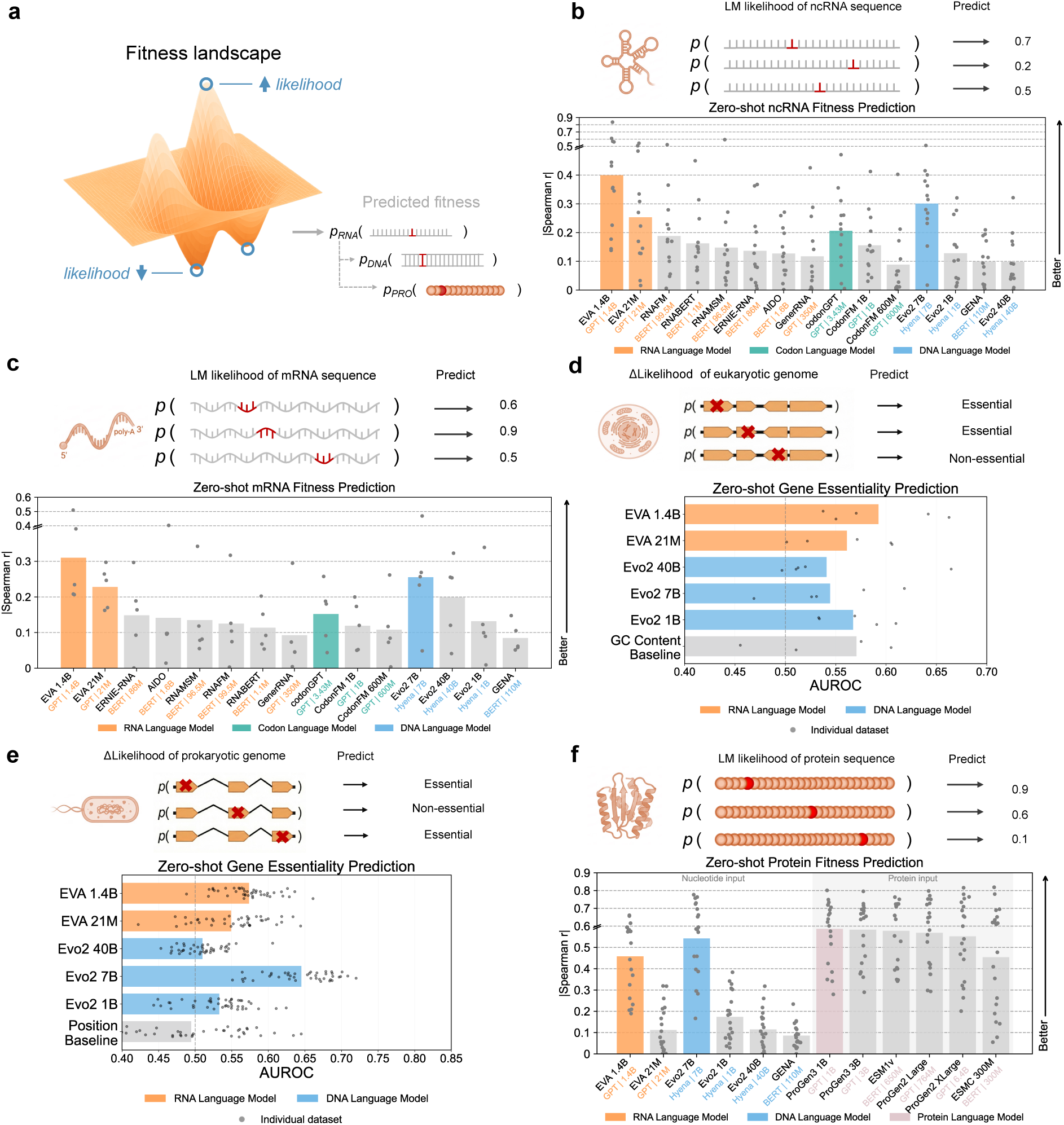
Zero-shot fitness prediction across RNA, DNA, and protein. (a) Conceptual overview of using EVA’s zero-shot sequence likelihood to approximate fitness landscapes and mutation effects for RNA, gene regions, and proteins. (b)(c) Benchmark results for zero-shot fitness prediction on ncRNA (b) and mRNA (c) sequences, evaluating the correlation between EVA likelihood scores and experimentally measured fitness across a diverse collection of deep mutational scanning (DMS) assays. Performance is reported as Spearman’s rank correlation (*ρ*); points indicate individual assays. (d)(e) Benchmark results on eukaryotic (d) and prokaryotic (e) gene essentiality prediction by inserting premature stop codons near the 5^′^ end of coding sequences and quantifying the EVA log-likelihood drop to infer essentiality. (f) Benchmark results for zero-shot fitness prediction on protein, evaluating the correlation between likelihood scores and experimentally measured fitness. Nucleotide-input EVA predictions are compared against protein language model baselines.

Since RNA is transcribed from DNA, EVA can also model mutational perturbations in gene regions. We formulate gene essentiality prediction as a likelihood-change assay, whereby five tandem premature stop codons are inserted near the 5^′^ end of each coding sequence, and the resulting drop in EVA log-likelihood relative to wildtype is used to infer whether the disrupted gene is functionally essential. On eukaryotic benchmarks, EVA surpasses Evo 7B and Evo2 40B despite being trained solely on RNA; on prokaryotic benchmarks, EVA’s 21M-parameter model even outperforms Evo2 40B and 1B (Figures 2D-E). These results reveal that transcriptomic sequences inherently encode the evolutionary signatures of gene functionality, allowing EVA to infer gene essentiality without direct exposure to genomic DNA.

Although EVA is trained exclusively on RNA, coding sequences directly determine the encoded protein, enabling EVA to capture evolutionary constraints relevant to protein function. On protein DMS benchmarks, EVA likelihood-based scoring is computed by conditioning on the mRNA type token, achieving a mean Spearman correlation of 0.46 and matching or exceeding specialized protein language models such as ESMC 300M on several datasets (Figure 2F). These results suggest that coding RNA sequences carry evolutionary fitness constraints that partially overlap with those learned by protein-centric models, offering preliminary evidence that RNA can serve as a cross-modal representation bridging transcriptomic and proteomic fitness landscapes.

### 2.3 EVA supports controllable RNA generation via CLM and GLM

EVA supports two complementary generation modes, Causal Language Modeling (CLM) for autoregressive sequence generation and Generalized Language Modeling (GLM) for masked infilling, both of which can be used with or without RNA-type and taxonomic conditioning tokens, enabling a spectrum from unconstrained sampling to fine-grained controllable design.

To quantify generation quality, we apply a lightweight filter to remove invalid samples based on GC content (30–70%), low-repeat content (*<*10%), and proper termination (see Methods). Incorporating RNA-type tags at pretraining markedly improves sample validity: the GC content pass rate increases from 47% without RNA-type pretraining (*n* = 2,000) to 97% for unconditional generation (*n* = 982) and 96% for conditional generation (*n* = 3,625), while repeat and termination filters reach near-perfect pass rates (≥99%) across all conditioned settings (Figure S5). These results demonstrate that RNA-type conditioning at pretraining improves the quality and biological validity of generated sequences.

We compare CLM and GLM in an unconditioned setting (Figure 3A). CLM spans both high-similarity and more divergent sequences relative to natural neighbours, reflecting the model’s ability to both reproduce and extend the natural manifold; GLM achieves higher reconstruction similarity by leveraging richer flanking sequence context. When conditioned on RNA-type tokens, GLM infilling achieves median reconstruction similarity of around 0.5 across all 11 RNA classes (Figure 3B), demonstrating that both modes benefit from structured conditioning. Unlike conventional RNA language models that learn an unstructured global distribution, EVA steers sampling toward specific RNA classes and host organisms–generalizing class-conditioned generation^24^ to RNA across all 11 major RNA classes.

**Figure 3:**
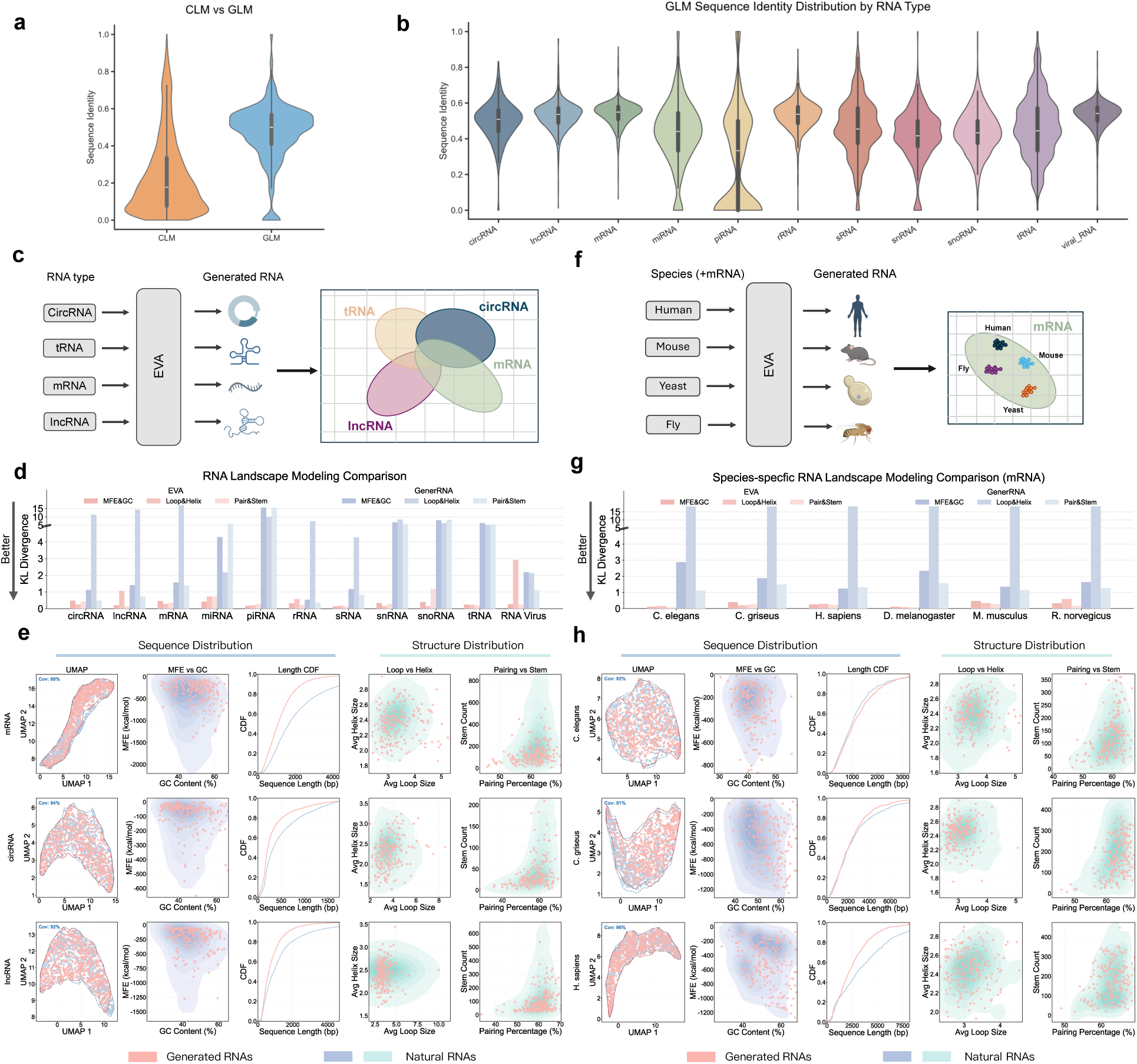
Controllable RNA generation with EVA across diverse RNA families and species. (a) Sequence similarity between generated sequences and their closest natural neighbours for unconditional CLM, and reconstruction similarity for GLM; CLM produces both high-similarity and more divergent sequences, whereas GLM achieves higher reconstruction via richer flanking context. (b) GLM infilling similarity between generated and masked fragments across 11 RNA classes. (c) Schematic of conditional generation controlled by RNA-type tags. (d) KL divergence comparison between EVA and GenerRNA across 11 RNA classes, evaluated over three secondary-structure feature groups (GC content & MFE, loop & helix composition, and base-pairing & stem statistics); EVA achieves *>*13-fold lower KL divergence. (e) Distributional comparisons of RNA-type conditioned samples vs. natural RNAs across UMAP embedding, length, MFE, GC content, and secondary-structure features. (f) Schematic of conditional generation controlled by taxonomic (species) tags. (g) KL divergence comparison between EVA and GenerRNA across six representative mRNA species; EVA achieves *>*30-fold lower KL divergence, reflecting accurate modeling of organism-specific RNA landscapes. (h) Distributional comparisons of species-conditioned mRNAs vs. natural sequences across sequence, biophysical, and structural metrics; extended results in Figure S7.

To benchmark generation quality, we compare EVA with GenerRNA ^15^—currently the only open-source RNA generative model; RNAGenesis ^16^ has not released model checkpoints and thus precludes direct comparison. Computing KL divergence on three secondary-structure feature groups (GC content & MFE, loop & helix composition, and base-pairing & stem statistics) across all 11 RNA classes, EVA achieves more than 13-fold lower KL divergence than GenerRNA, with particularly large improvements for piRNA, snRNA, snoRNA, and tRNA (Figure 3D; Supplementary Table S11). We further compare generated sequences with natural RNAs across sequence-space and biophysical statistics, including UMAP embedding, length distribution, MFE, GC content, and secondary-structure features; RNA-type conditioning yields closer agreement with natural distributions than unconditional generation (Figure 3E). This reflects the model’s ability to capture family-specific sequence and structural grammars—for example, the high GC content and stable folds of rRNAs, or the short length and minimal structure of miRNAs. Extended distributional comparisons across all 11 RNA classes are provided in Figure S6.

After fixing RNA type (e.g., mRNA), we further condition generation on species tags. Restricting to mRNA, EVA achieves more than 30-fold lower KL divergence than GenerRNA across six representative species (*C. elegans*, *C. griseus*, *H. sapiens*, *D. melanogaster*, *M. musculus*, and *R. norvegicus*), demonstrating that species-aware conditioning enables accurate modeling of organism-specific RNA sequence and structural landscapes (Figure 3G; Supplementary Table S12). Distributional comparisons across sequence and structural metrics confirm that species conditioning further improves agreement between generated and natural sequences across *C. elegans*, *C. griseus*, and *H. sapiens* (Figure 3H). Extended results across additional RNA classes and species are provided in Figure S7.

### 2.4 EVA generates novel RNAs with structural fidelity and extends to unseen RNA classes via fine-tuning

Having established that EVA’s conditional generation recapitulates natural sequence and secondary structure distributions, we next examine whether the model also captures tertiary structural information. tRNA is an informative test case, as its L-shaped fold is structurally well-defined, functionally critical, and must be encoded entirely in sequence.

Structural similarity was assessed by TM-score between AlphaFold3-predicted structures and all experimentally resolved tRNA structures from RCSB; sequence novelty was quantified as maximum sequence identity against 10,000 randomly sampled tRNAs from training data. Across the generated ensemble, structural similarity to natural tRNAs and sequence divergence from known tRNAs are largely decoupled (Figure 4A), with generated sequences maintaining a mean TM-score of 0.47 even as sequence identity to training tRNAs drops to approximately 0.55, consistent with EVA capturing fold-level constraints independently of specific sequences. Structural alignment of selected generated sequences against their closest experimentally resolved counterparts in RCSB illustrates this at the level of individual examples (Figure 4B), where representative cases achieve a mean TM-score of approximately 0.74 at an average sequence identity of only 0.45, demonstrating that overall backbone topology is preserved despite substantial sequence divergence from any known tRNA. Together, these observations suggest that training on evolutionarily diverse sequences implicitly encodes fold-level structural constraints, consistent with the notion that EVA’s learned distribution approximates an evolution-shaped manifold in which sequence novelty and structural viability coexist.

**Figure 4:**
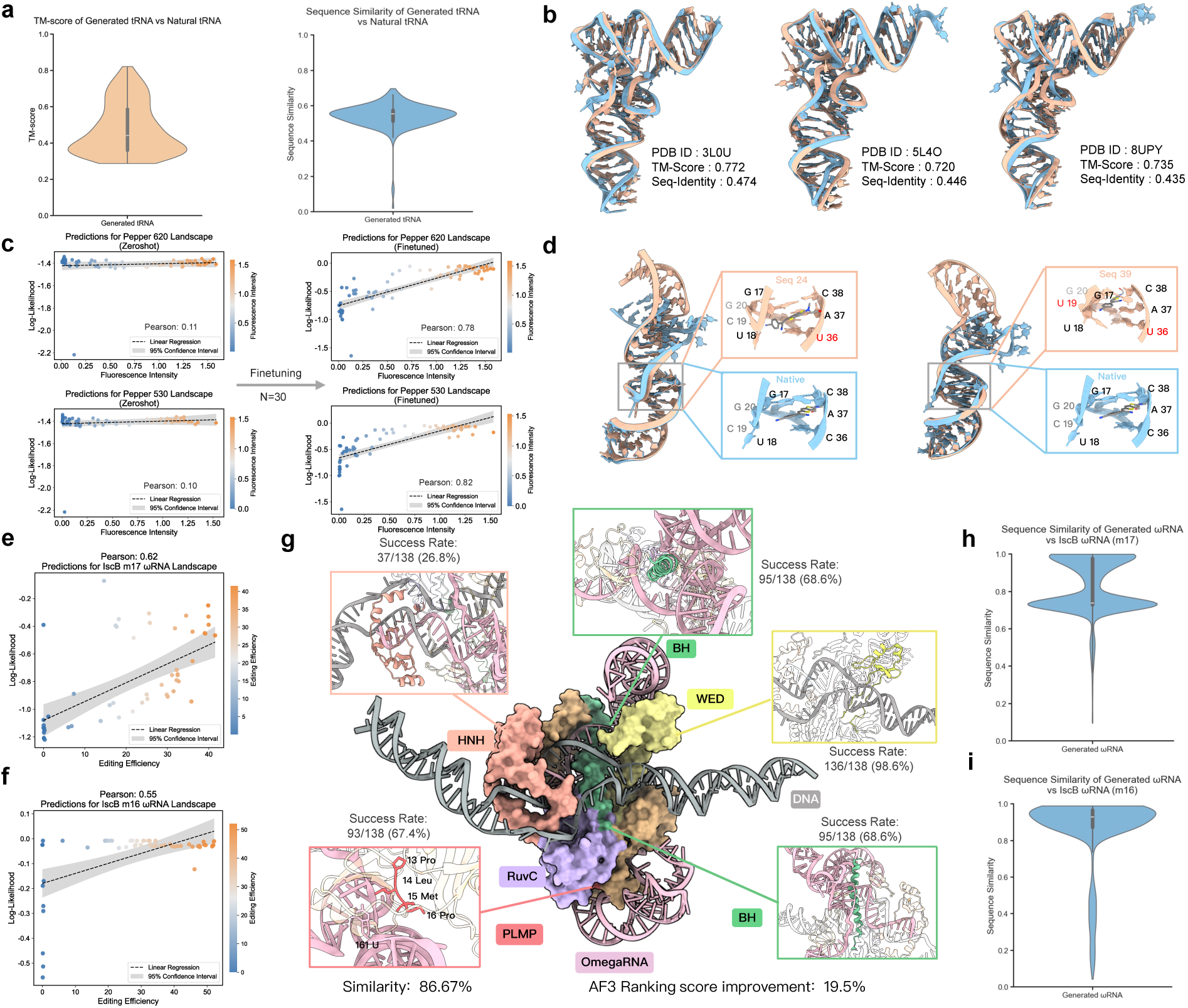
De novo design of tRNAs, RNA aptamers, and CRISPR gRNAs. (a) Zero-shot de novo generation of tRNAs showing high structural similarity to natural tRNAs, quantified by TM-score, while maintaining sequence novelty relative to known tRNAs. (b) Representative structural alignments between AlphaFold3-predicted structures of generated tRNAs and experimentally resolved natural tRNAs, illustrating preservation of the canonical L-shaped fold despite low sequence identity. (c) Prediction of fluorescence intensity for RNA aptamers after low-data fine-tuning (n = 30). Left: binding to HBC530; right: binding to HBC620. Model log-likelihood scores are compared against experimentally measured fluorescence. (d) Structural comparison between de novo generated aptamers (blue) and wildtype structures, showing preservation of the ligand-binding pocket and overall fold despite sequence mutations, consistent with learned mutational-structural relationships. (e,f) Prediction of editing efficiency for IscB omegaRNAs after fine-tuning on m16 (n = 42) and m17 (n = 44) sequence sets, respectively. (g) AlphaFold3-predicted complex structure of IscB protein, de novo designed m16 omegaRNA, and target DNA. Success is defined by satisfying a set of domain-level structural criteria relative to wildtype (see Methods). (h,i) Violin plots showing sequence similarity distributions between de novo generated omegaRNAs and the corresponding wildtype sequences for m17 (h) and m16 (i). Generated sequences span a biologically reasonable range of divergence while remaining structurally plausible.

While EVA supports conditional generation for 11 major RNA classes, many valuable RNAs remain outside this scope. RNA aptamers are synthetically engineered molecules absent from nature, and guide RNAs from metagenomic CRISPR-Cas systems represent highly specialized, application-specific sequences. To enable design beyond pre-trained families, we reserved a blank conditioning token to support fine-tuning and conditional generation for user-defined RNA types.

We fine-tuned EVA on a small set of high-fluorescence aptamer sequences (*n* = 30), using sequence-level self-supervised language modeling to steer the model’s generative distribution toward the functional aptamer sequence space. After fine-tuning, EVA log-likelihood scores correlate with experimentally measured fluorescence intensity for aptamers binding to both HBC530 and HBC620 ligands, with Spearman correlation improving from 0.1 (zero-shot) to 0.8 (Figure 4C). Position-wise comparison of mutation enrichment confirms that the fine-tuned model preferentially generates the same nucleotide substitutions that experimentally confer improved fluorescence activity, as revealed by alignment with the DMS fitness landscape (Figure S8A, left). Generated aptamers show mutation counts within a biologically reasonable range relative to wildtype (Figure S8A, right). De novo generated aptamers show preserved ligand-binding pocket geometry despite sequence mutations in predicted structures (Figure 4D), providing an in silico indication of structural plausibility.

IscB is an ancestral CRISPR-associated transposon protein that pairs a compact protein with a large guide RNA, the *ω*RNA^25^. A central engineering challenge is to reduce *ω*RNA size while preserving target-site recognition, enabling more efficient viral delivery. We applied the same fine-tuning strategy using high-editing-efficiency sequences from two guide RNA variants. After fine-tuning, EVA log-likelihood correlates with editing efficiency for both variants (Figure 4E, 4F), and more than half of de novo generated sequences are shorter than the respective wildtype (Figure S8B). Structural plausibility was assessed using five domain-level geometric criteria derived from published IscB structural analyses ^25,26^ (Table S13); designs satisfying all criteria are flagged as in silico candidates for experimental validation (Figure 4G). Generated *ω*RNAs span a biologically reasonable range of sequence identities relative to wildtype (Figure 4H, 4I). These results demonstrate that EVA adapts to RNA classes absent from its training distribution using minimal data.

### 2.5 EVA enables mRNA and circRNA vaccine design

We next assess whether EVA can be used as a practical design engine for RNA vaccines. RNA vaccines have emerged as a transformative therapeutic modality, yet their design remains challenging as efficacy depends on multiple sequence-level properties–codon usage, folding stability, and translational efficiency–that must be jointly optimized. Circular RNAs have attracted increasing interest as vaccine platforms in recent years, owing to their covalently closed topology, which confers markedly greater resistance to exonucleolytic degradation compared to linear mRNA and can extend the duration of antigen expression in vivo^27,28^. However, circRNA vaccine design presents an additional engineering challenge beyond ORF codon optimization. Because the molecule lacks a 5^′^ cap and poly(A) tail, translation must be entirely driven by an internal ribosome entry site (IRES), making IRES design a critical determinant of translational efficacy ^27,28^.

The standard pipeline for mRNA codon optimization jointly optimizes minimum free energy (MFE) for RNA folding stability and the Codon Adaptation Index (CAI) for host codon usage compatibility ^29^. However, CAI treats each codon position independently and relies on fixed frequency tables, lacking the ability to model broader sequence context or organism-specific translational dynamics ^30–36^. We therefore replace CAI with EVA log-likelihood under joint RNA-type and species conditioning, which captures host-organism compatibility in a context-aware, unified score while retaining MFE as the folding objective (Figure 5A). We perform in silico codon optimization on five mRNA vaccine systems (SARS-CoV-2 Spike, Influenza PR8 HA (A/PR/8/34 H1N1), Varicella-Zoster Virus (VZV) gE, RABV-G, and HIV) and benchmark against Evo2-1B ^19^ and CodonFM-1B ^32^ as baselines. EVA-guided synonymous search improves MFE while simultaneously increasing CAI compared to both baselines; joint RNA-type and species conditioning further boosts performance by aligning the optimization trajectory with the class- and organism-specific sub-manifold (Figure 5B). Detailed numerical results for all five vaccine systems and all baselines are provided in Supplementary Table S5.

**Figure 5:**
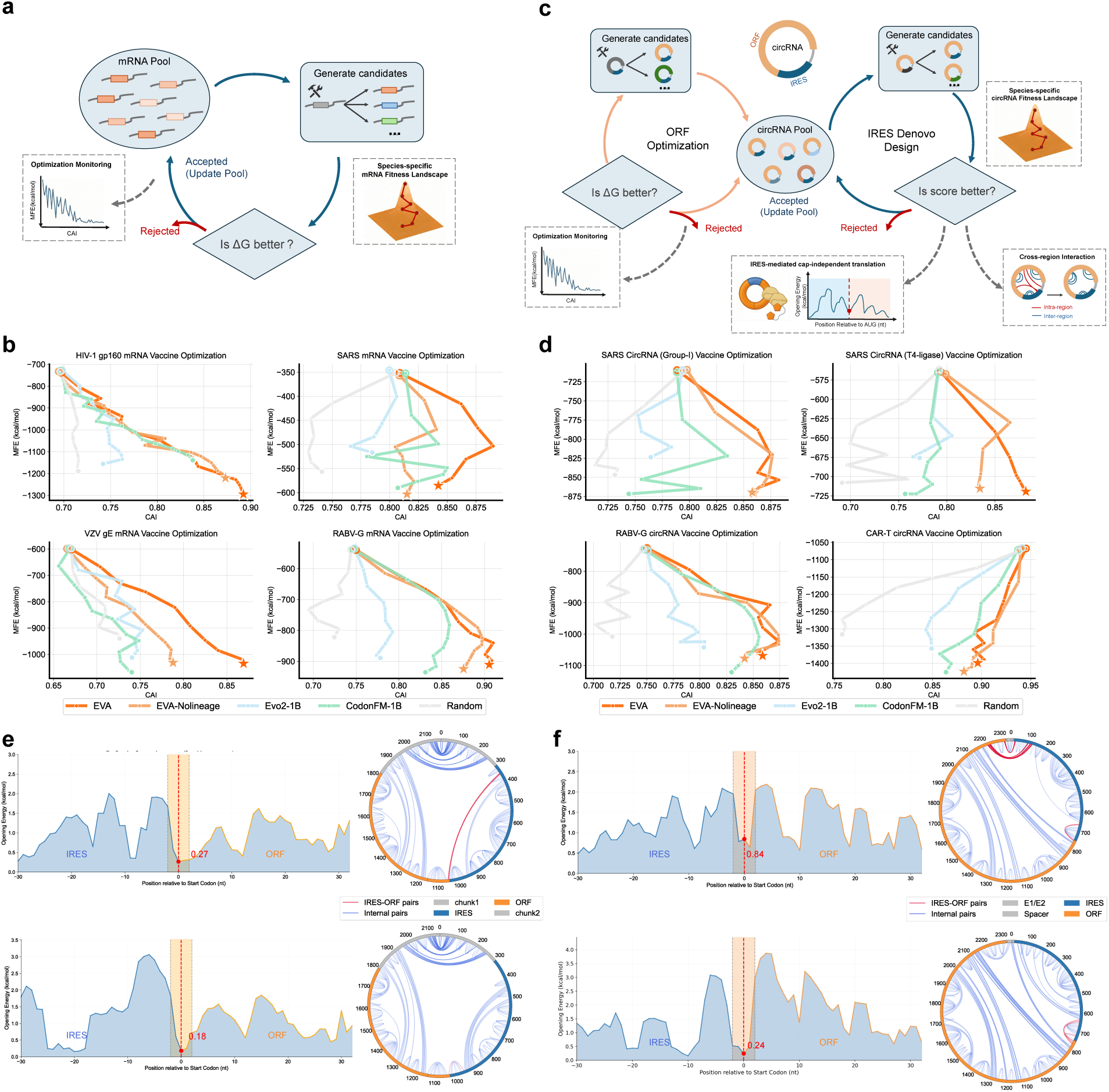
mRNA and circRNA vaccine design with EVA. (a) Conceptual overview of mRNA codon optimization with EVA. (b) Benchmark on five mRNA vaccine systems (SARS-CoV-2 Spike, Influenza PR8 HA, VZV gE, RABV-G, HIV) comparing CDS optimization in terms of MFE and CAI against Evo2-1B, CodonFM-1B, and a Random (MFE-only) baseline, all conditioned on the mRNA type and *Homo sapiens*. (c) Conceptual overview of circRNA vaccine design with EVA: ORF codon optimization via species-conditioned autoregressive generation (left), and de novo IRES redesign via EVA GLM where the IRES region is masked and regenerated conditioned on flanking sequences (right). (d) Benchmark on four circRNA vaccine systems (SARS: Group-I, SARS: T4 ligase, rabies, CAR-T) for ORF codon optimization, same baselines and species conditioning as (b). (e)(f) Biophysical properties of EVA-redesigned IRES sequences versus the original CVB3 IRES for SARS and rabies circRNA vaccines, including ribosome accessibility and inhibitory long-range interactions.

We apply the same EVA-guided codon optimization pipeline (MFE + EVA log-likelihood, conditioned on the circRNA type and *Homo sapiens*) to circRNA ORF optimization (Figure 5C). We evaluate four circRNA systems (SARS: Group-I, SARS: T4 ligase, rabies, and CAR-T) against the same baselines (Evo2-1B^19^ and CodonFM-1B ^32^) and observe consistent improvements in both CAI and MFE, with additional gains when conditioning on the target species (Figure 5D). Detailed ORF optimization results for all circRNA systems are provided in Supplementary Table S6.

Beyond ORF optimization, circRNA translation depends entirely on the Internal Ribosome Entry Site (IRES). De novo IRES design offers the potential for reduced immunogenicity, improved circularization compatibility, and enhanced cell-type specificity compared to fixed viral elements such as CVB3^27^.

We address this by leveraging EVA’s GLM capability, masking the IRES region and conditioning generation on the flanking ORF and cyclization sequences (circRNA type, *Homo sapiens*). This enables the model to perform de novo design of IRES within its molecular context (Figure 5C). Designed IRES candidates were evaluated on two key biophysical criteria (ribosome accessibility and inhibitory long-range interactions; see Methods). Across both SARS and rabies circRNA systems, EVA-generated IRES candidates outperform the original CVB3 IRES across both biophysical metrics. For SARS circRNA, inhibitory long-range interactions are reduced by 58% and ribosome accessibility is increased by 33% (Figure 5E). For rabies circRNA, inhibitory long-range interactions are reduced by 27% and ribosome accessibility is increased by 71% (Figure 5F).

### 2.6 Feature interpretation in EVA from neurons to biological properties

Despite EVA’s strong performance across prediction and generation tasks, it remains important to ask what the model has actually learned from RNA sequences. Large language models are often characterized as opaque, yet mechanistic interpretability methods offer tools to examine their internal representations directly. We apply sparse autoencoders (SAEs) ^37^ to decompose the dense, distributed activations of EVA’s residual stream into a high-dimensional set of sparse features, each of which can be examined for biological correspondence (Figure 6A). SAEs were trained on layer-13 activations extracted from one million RNA sequences spanning diverse RNA types and taxonomic groups (see Methods).

**Figure 6:**
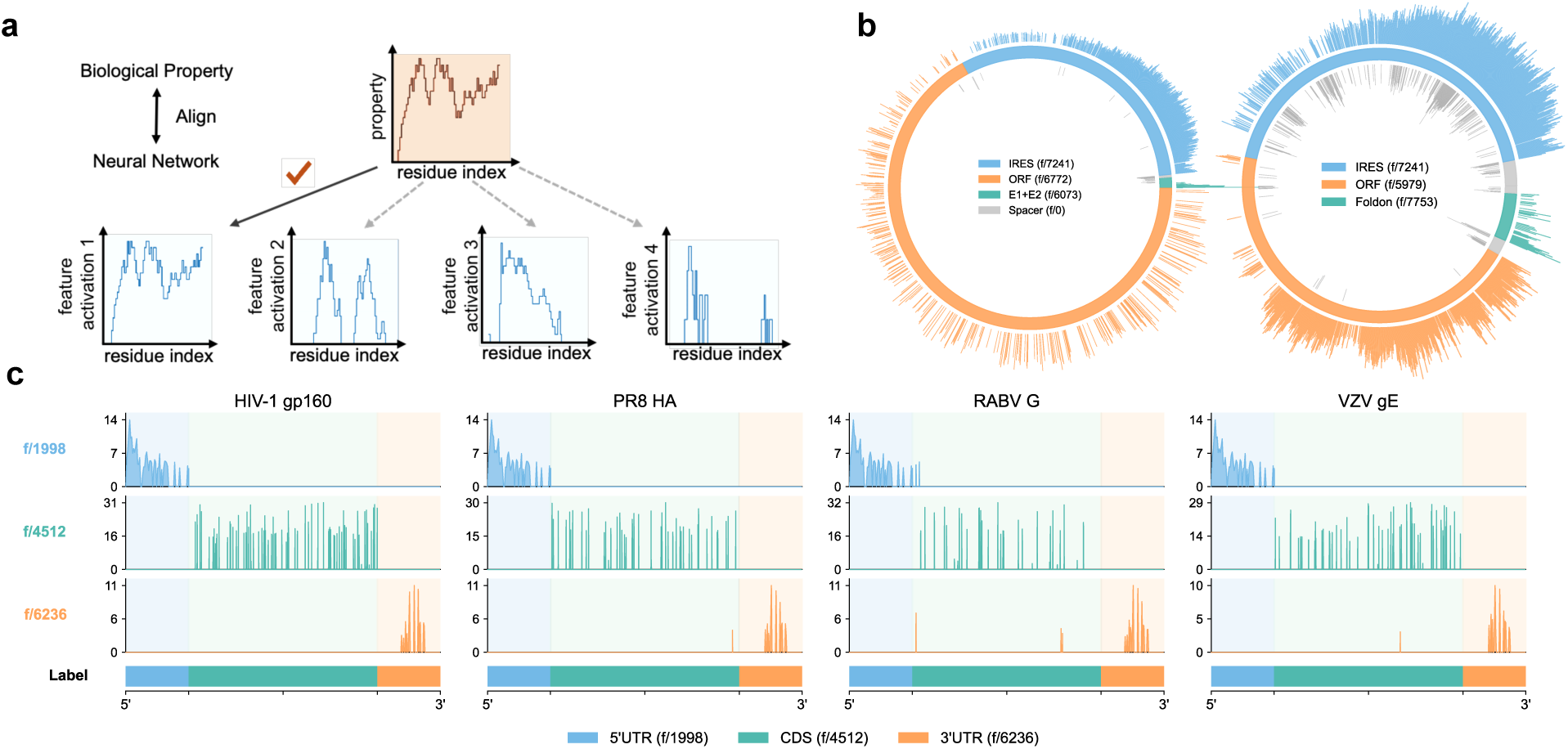
Interpretability analysis revealing hierarchical RNA features learned by EVA. (a) Schematic illustrating how model neurons are mapped to biological concepts using sparse autoencoders, revealing that EVA can learn structure- and function-related biological knowledge from sequence alone. (b) Activations of features associated with IRES, ORF, and related functional elements in circRNA vaccines (left: RABV-G; right: SARS-CoV-2). (c) Activations of features associated with 5^′^UTR, CDS, and 3^′^UTR in mRNA vaccines (HIV, PR8 HA, RABV-G, and VZV gE).

To assess whether learned features correspond to known RNA biology, we identify features whose activation is systematically enriched over specific functional sequence elements. In circRNA vaccine sequences (RABV-G and SARS-CoV-2), individual SAE features activate selectively over IRES and ORF regions (Figure 6B), suggesting that the model has formed internal representations that track these regulatory and coding boundaries. In mRNA vaccine sequences (HIV, PR8 HA, RABV-G, and VZV gE), three distinct features consistently demarcate the 5^′^UTR, CDS, and 3^′^UTR across all four constructs, despite substantial differences in their coding sequences (Figure 6C). This cross-construct consistency indicates that these features reflect generalizable positional and functional logic rather than sequence-specific memorization, suggesting that EVA has learned not only the identity of individual functional elements but also their characteristic spatial arrangement within a transcript.

At a higher organizational level, SAE features cluster according to the biological domain of origin: features preferentially activated by eukaryotic sequences and those preferentially activated by prokaryotic sequences occupy spatially separated regions in the UMAP projection of SAE decoder directions (Figure S10). Individual experts within EVA’s Mixture-of-Experts backbone further exhibit non-uniform activation profiles across RNA types (Figure S10), consistent with a degree of functional specialization emerging from training. Together, these observations suggest that EVA’s internal representations are organized along biologically meaningful axes, from domain-level taxonomy to transcript-level functional architecture.

## 3 Discussion

EVA demonstrates that large-scale generative learning on evolutionary RNA sequence diversity yields a unified framework supporting both sequence scoring and controllable design. By training on 114 million full-length RNA sequences with structured conditioning on RNA type and taxonomy, EVA learns an evolution-consistent representation in which sequence likelihood reflects evolutionary plausibility, enabling zero-shot mutation-effect prediction across RNA, DNA gene regions, and proteins without task-specific retraining. Conditioned generation across 11 RNA classes, enabled in part by an 8,192-token context window that accommodates full-length transcripts beyond the reach of prior models, produces sequences that recapitulate natural length, folding energy, and structural distributions while remaining evolutionarily novel, and practical applications spanning tRNA design, aptamer engineering, CRISPR guide RNA generation, and mRNA/circRNA vaccine optimization demonstrate the breadth of this capability.

A notable finding is that, enabled by EVA’s mixture-of-experts architecture and joint RNA-type and taxonomic conditioning that disentangles the heterogeneous RNA sequence landscape, the learned representations are interpretable and structurally organized. Sparse autoencoder analysis reveals that individual features localize to functional elements, including 5^′^UTRs, CDSs, 3^′^UTRs, IRESs, and ORFs, in a manner that generalizes across diverse constructs, indicating that EVA has learned the functional architecture of RNA from sequence alone. Complementarily, experts in the MoE backbone exhibit RNA-type-specific activation patterns, and features segregate by biological domain in the SAE latent space, together suggesting that EVA’s internal organization mirrors the hierarchical structure of RNA biology.

The current work is limited to computational evaluation; wet-lab validation of designed sequences is ongoing. EVA also operates at the sequence level and does not explicitly model three-dimensional structure, which may limit applicability to tasks requiring precise conformational control. Additionally, the granularity of RNA type classification warrants further refinement, and the range of RNA classes supported for controllable generation without fine-tuning remains an area for expansion.

Looking forward, we see two directions of particular interest. First, scaling the model architecture and extending the context window could enable design at the level of complete RNA systems, moving from individual functional elements toward more complex assemblies such as RNA viral genomes; recent work demonstrating the generative design of viable bacteriophages from genome language models provides an encouraging proof of concept ^38^. Second, the interpretability analyses presented here establish a foundation for a complementary goal: not merely identifying which internal features correspond to biological concepts, but actively steering those features to guide generation toward desired functional outcomes^39–41^. Together, these directions point toward generative models that are both more capable and more controllable, advancing the generative design of increasingly complex RNA systems as a step toward generative biology.

## 4 Methods

### 4.1 Model Training

EVA is trained on RNA sequences from OpenRNA v1 using a two-stage curriculum that combines Causal Language Modeling (CLM) and Generalized Language Modeling (GLM) objectives throughout both stages, while progressively introducing richer conditioning information. In Stage 1 (pre-training), the model is conditioned on RNA-type tags and learns universal RNA sequence grammar. In Stage 2 (mid-training), taxonomic lineage tags are additionally introduced, enabling species-aware generation. We trained EVA across a range of model scales on 16 NVIDIA H200 141 GB GPUs across 2 nodes, and observe consistent perplexity reduction with increasing capacity. Benchmarks and evaluations are conducted on the smallest (~21M parameters) and largest (1.4B parameters) checkpoints; all intermediate checkpoints are publicly released. All experiments use a context length of 8,192 tokens.

#### 4.1.1 Model Architecture

EVA is a decoder-only Transformer with an 8,192-token context window. It incorporates rotary positional embeddings (RoPE, *θ* = 10^5^), multi-head attention (MHA), and pre-layer RMSNorm normalization. Feed-forward layers are replaced by a Mixture-of-Experts (MoE) backbone implemented via MegaBlocks, with 8 experts per layer of which 2 are activated per token (top-2 routing) and a capacity factor of 1.5. An auxiliary load-balancing loss with coefficient *λ*_aux_ = 0.01 is added to prevent expert collapse. Full architecture configurations and training hyperparameters for all four model scales are reported in Supplementary Table S4.

#### 4.1.2 Training Objectives

##### Mixed CLM and GLM pre-training

During training, each example is assigned one of two objectives sampled per sequence: (i)Causal Language Modeling (CLM), in which the model performs standard autoregressive next-token prediction over the full sequence; or (ii)Generalized Language Modeling (GLM), in which one or more contiguous spans are replaced by unique sentinel tokens (`<span_n>`) and the model predicts the masked tokens conditioned on the surrounding context and the corresponding sentinel, enabling fill-in-the-middle infilling. The GLM infilling objective generalizes the fill-in-the-middle (FIM) paradigm originally introduced for code language models ^42,43^ and subsequently adopted by protein language models such as xTrimoPGLM ^44^ and ProGen3^21^; we adapt this technique to RNA sequence modeling. Following ablation results from ProGen3^21^, we use a CLM:GLM mixing ratio of 2:1 (*p*_GLM_ = 0.333). For GLM examples, the total coverage ratio of masked tokens is drawn from 0.15, 0.25, 0.50, 0.80 with probabilities (0.28, 0.30, 0.28, 0.14); individual span lengths follow a mixture of Gaussian distributions (mean/std: 10/5, 20/10, 50/20 tokens) with mixing weights (0.3, 0.5, 0.2), capped at 10 spans per sequence and with no overlap between spans.

The composite training loss is:

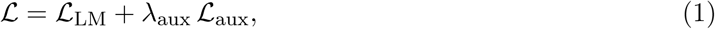

The language modeling loss 𝓛_LM_ is the mean cross-entropy over all non-padding target positions 𝒮 (positions where the label is not the ignore index −100):

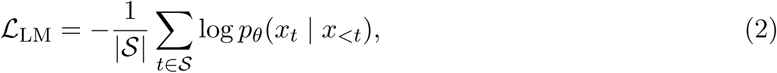

where for CLM examples 𝒮 spans all sequence tokens, and for GLM examples 𝒮 contains only the masked span tokens (all other positions carry label 100 and are excluded from the loss).

The auxiliary load-balancing loss 𝓛_aux_ follows the Switch Transformer formulation ^45^, encouraging uniform expert utilization across the batch:

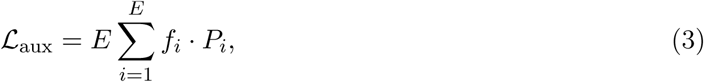

where *E* is the number of experts, 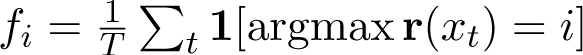 is the fraction of tokens routed to expert *i*, 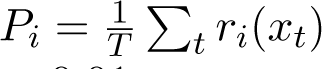 is the mean router probability for expert *i* over all *T* tokens in the batch, and *λ*_aux_ = 0.01.

##### Position-Preserving Fuzzy Encoding

For the GLM infilling objective, we adopt a position-preserving fuzzy encoding scheme. Given an RNA sequence with one or more masked spans, each span is replaced in-place by a unique sentinel token (`<span_n>`, *N* ϵ {0, . . . , 49}); the resulting context (with sentinels) is presented to the model first, terminated by `<eos>`, after which each span’s masked content is appended sequentially—each delimited by its corresponding sentinel and a closing `<eos_span>` token.

Position IDs are assigned as follows. In the *context* segment, tokens before the first masked span retain their original sequence positions; each sentinel `<span_n>` receives the position ID of the first masked nucleotide of its span (adjusted by any previously accumulated offset); and all context tokens that follow the *k*-th span have a cumulative *fuzzy offset* 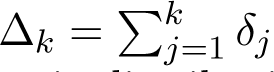 added to their original positions, where each *δ_j_*is sampled independently from a geometric distribution with mean 0.2 × *L_j_* (*L_j_* = length of span *j*). In the *infilling* segment (after `<eos>`), each sentinel is placed at the *original* (un-offset) position of the first masked nucleotide; the masked tokens that follow receive positions *p*_start_ + 1*, p*_start_+ 2*, . . .* (where *p*_start_ is the sentinel position), and `<eos_span>` is placed one position after the last masked token. Because the fuzzy offsets make the positional gap between a sentinel and the subsequent context tokens ambiguous, the model cannot infer the exact length of the masked span from positional cues alone, and must instead learn to terminate infilling by generating the content-based `<eos_span>` token.

A concrete RNA example is shown below, with two masked spans highlighted. Conditioning prefix tokens (lineage tag, RNA-type tag) precede the shown sequence in actual training inputs but are omitted here for clarity.

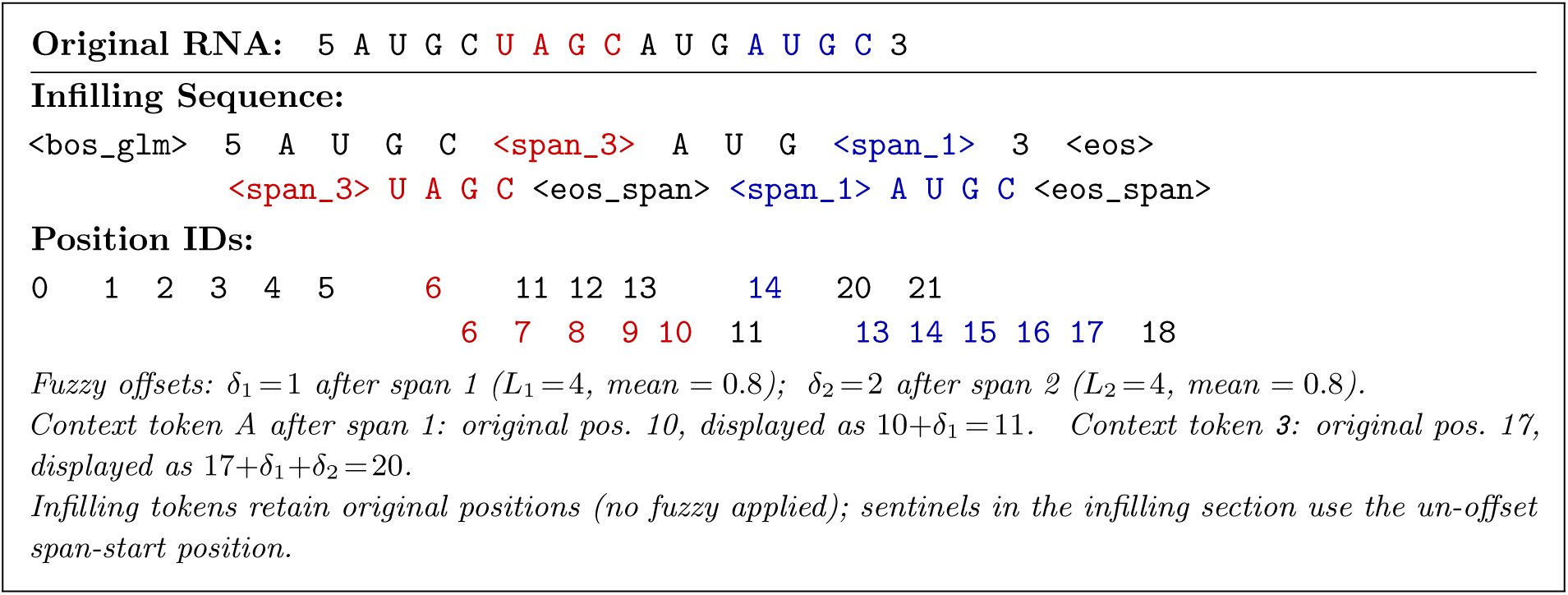

##### Input format and conditioning tokens

Each sequence is prepended with structured conditioning tokens that encode RNA-type identity (*e.g.*, `<rna_tRNA>`, `<rna_mRNA>`) and strand directionality (5 or 3). In Stage 2 (mid-training), a Greengenes-format taxonomic lineage tag is additionally prepended at the outermost level, resulting in inputs of the form:

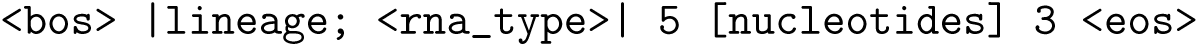

where the lineage string follows the seven-level Greengenes hierarchy (`d domain; p phylum; …; s species`). For GLM examples, `<bos>` is replaced by `<bos_glm>` and the sequence tokens contain the sentinels and infilling structure described above. Reverse-complement augmentation is applied with probability 0.5 in both stages: a sequence originally read 5 [seq] 3 is randomly swapped to 3 [`rev-comp`] 5, augmenting coverage of directional sequence patterns.

#### 4.1.3 Two-Stage Training Curriculum

##### Stage 1: RNA-type-conditioned Pre-training

In Stage 1, the model is trained on the full 114M-sequence OpenRNA v1 dataset conditioned only on RNA-type tags. The mixed CLM+GLM objective enables the model to simultaneously learn autoregressive generation and span infilling. By withholding fine-grained taxonomic information, Stage 1 focuses the model on universal RNA sequence grammar, including shared structural motifs, canonical base-pairing patterns, and family-level syntax, across all 15 RNA categories.

##### Stage 2: Lineage-conditioned Mid-training

In Stage 2, training resumes from the Stage 1 checkpoint, with only model weights loaded; the optimizer state and learning-rate scheduler are fully reset. The key change is the introduction of Greengenes-format phylogenetic lineage tags as additional conditioning prefixes. To prevent catastrophic forgetting of the universal RNA grammar learned in Stage 1, 20% of training examples are kept in the pre-training (no-lineage-prefix) format, while 80% use the full lineage-conditioned format. The mixed CLM+GLM objective is retained. This staged approach, which learns RNA-type-level grammar first and subsequently incorporates species-specific constraints, proves critical; empirically, introducing lineage conditioning from the outset leads to overfitting, whereas the two-stage curriculum achieves substantially lower perplexity and better generation quality (Figure S4B).

#### 4.1.4 Data Sampling Strategy

A major challenge in training RNA foundation models is the extreme imbalance and low evolutionary conservation of RNA sequences compared to proteins. The natural RNA landscape follows a heavy-tailed distribution, dominated by a few massive clusters of housekeeping RNAs (e.g., rRNAs, tRNAs) while containing numerous small clusters of diverse, functional non-coding RNAs. We found that direct training on this raw distribution causes the model to severely overfit to abundant clusters while neglecting rare families (Figure S3). To overcome this, we implemented an evolutionary conservation-based sampling strategy. Sequences were first clustered by similarity to estimate the evolutionary density of each family. During training, sampling probability was re-weighted in inverse proportion to the square root of cluster size, effectively down-sampling abundant housekeeping RNAs and up-sampling rare functional RNAs. This strategy flattens the training distribution, ensuring the model captures the full diversity of the RNA landscape. Comparative training dynamics confirm that this approach prevents overfitting to dominant clusters, including rRNA and tRNA, and yields substantially improved model generalization compared with uniform or inverse-linear sampling (Figure S2C, S2D).

#### 4.1.5 Optimization and Hyperparameters

All models are trained with the AdamW optimizer (*β*_1_ = 0.9, *β*_2_ = 0.95, *ε* = 10^−8^, weight decay = 5 × 10^−6^). The learning rate follows a linear warm-up schedule (3,000 steps in Stage 1; 1,000 steps in Stage 2), followed by cosine annealing to a minimum of 10% of the peak learning rate. Based on systematic hyperparameter search, we use a peak learning rate of 1 × 10^−4^ (Figure S4A). The effective batch size is 256 sequences (4 sequences per device × 16 gradient accumulation steps × 4 data-parallel replicas). All models are trained in BF16 mixed precision.

#### 4.1.6 Scaling Law Analysis

Scaling laws describe how model performance improves with compute and can guide optimal allocation of pre-training resources between model size and data^46,47^. To characterize EVA’s scaling behavior, we fit a Kaplan-style power law ^46^ to the final validation perplexities of four model scales (21M, 145M, 437M, and 1.4B parameters). Training FLOPs *F* are computed based on active parameters (i.e., the subset of parameters activated per forward pass in the MoE architecture), following standard practice for sparse models. We fit the power law in log space, modeling *L* = ln(PPL) as a function of *F* :

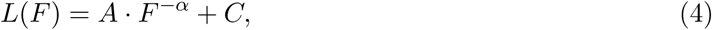

fitted via nonlinear least squares using `scipy.optimize.curve_fit`. The fitted parameters are *A* = 75867.99, *α* = 0.3211, and *C* = 1.2444, yielding a power-law decrease in perplexity (Figure 1I).

#### 4.1.7 Distributed Training Infrastructure

The 1.4B model employs a hybrid parallelism strategy across 16 H200 141 GB GPUs (2 nodes, 8 GPUs each). **Expert parallelism** (EP = 4) shards the 8 MoE experts across 4 GPUs via All-to-All communication using the MegaBlocks framework; each GPU holds 2 experts and processes tokens routed to its local experts. Orthogonally, **data parallelism** (DP = 4) replicates each EP group across 4 independent data-parallel ranks, with gradients of non-expert parameters synchronized via All-Reduce across the DP group after each gradient-accumulation cycle. The resulting EP DP = 4 4 = 16 GPU layout achieves near-linear throughput scaling while keeping per-device peak memory within the 141 GB limit.

### 4.2 OpenRNA v1 training data

Existing large-scale nucleotide databases such as MARS ^17^ aggregate extensive sequence collections but intermix RNA and DNA data; models trained on such mixed-modality corpora have been shown to underperform relative to those trained on curated, modality-specific data^10^, motivating the construction of a dedicated, high-quality, full-length RNA corpus. To build OpenRNA v1, we surveyed 19 public RNA resources and integrated 15 databases into a unified atlas containing 114M curated full-length RNA sequences. More than 90% of sequences are annotated with both RNA type and taxonomic labels. OpenRNA v1 covers 15 RNA categories, including mRNA, lncRNA, circRNA, snoRNA, snRNA, sRNA, tRNA, rRNA, piRNA, miRNA, YRNA, scaRNA, ribozyme, viral RNA and vaultRNA.

#### 4.2.1 Data curation

OpenRNA v1 was built by integrating sequences from eight primary sources: NCBI Nucleotide and Virus databases (NT & Virus; 56.2M sequences after filtering, 49.25%), RNACentral Consortium aggregating Rfam, GtRNAdb and partner databases (33.4M, 29.25%), Ensembl vertebrate genomes Release 114 (15.7M, 13.75%), three merged circRNA databases (circBase, circAtlas; 2.1M, 1.87%), SILVA high-quality ribosomal RNA alignments (0.55M, 0.48%), NONCODE long non-coding RNAs (0.22M, 0.19%), piRNAdb (42K, 0.04%), and a collection of organism-specific databases including WormBase, FlyBase, snoDB, and miRBase (5.9M, 5.17%). Sequences were first filtered to remove redundant entries arising from cross-database overlap, low-complexity sequences (e.g., homopolymer-dominated transcripts), and sequences with ambiguous nucleotide content exceeding 5%. Following filtering, 114M unique full-length RNA sequences were retained (Supplementary Table S2).

##### Core and Genomic Archives

We utilized RNACentral and Rfam as the structural backbone. For Rfam, we implemented a dual-resolution labeling strategy: sequences with definitive family-level evidence were assigned specific functional labels (e.g., *snRNA*, *snoRNA*), while transcripts lacking high-resolution classification but exhibiting characteristic short-RNA length profiles (50–200 nt) were curated as *sRNA*. This category serves as a reservoir of structural candidates. To ensure broad taxonomic coverage, we integrated Ensembl (341 species) and the NCBI Nucleotide (NT) database. For Ensembl, we filtered out low-complexity regions (e.g., poly-A/T stretches). For the massive NCBI NT collection, we employed `accession2taxid` for precise species mapping and removed sequences with ambiguous taxonomic lineage.

##### Specialized Functional RNA Enrichment

We enriched the dataset with specialized repositories to capture distinct RNA modalities. For lncRNAs, we integrated NONCODE, LNCipedia, LncRNAWiki, and LncRNADisease; poly(A) tails in LNCipedia entries were computationally trimmed to avoid homopolymer bias. For small RNAs, miRBase and MirGeneDB were unified for miRNAs, explicitly distinguishing mature sequences from precursors and excluding incomplete stem-loop fragments; piRNAdb and snoDB were incorporated to cover piRNAs and snoRNAs, respectively. circRNA sequences (from circBase) were standardized to linear representations for tokenization. rRNA sequences were drawn from SILVA and PR2, with primer sequences, vector contamination, and potential chimeras removed from SILVA entries. Viral RNA coverage was established using NCBI Virus, classifying sequences into six viral RNA types (e.g., dsRNA, ssRNA+, ssRNA) and filtering fragmented contigs shorter than 20 nt.

##### Deduplication and Filtering Pipeline

We developed a unified key-value header format to resolve schema discrepancies across the 15 databases. Redundancy was addressed in two stages: intra-database exact duplicates were removed based on SHA256 content hashing, followed by inter-database deduplication using `seqkit` (v2.8.2) with the `rmdup –by-seq` option and the xxHash algorithm for high-efficiency cross-corpus sequence comparison.

Finally, we applied a strict functional filter to retain 15 core RNA categories (e.g., mRNA, rRNA, tRNA, lncRNA, sRNA, viral RNA). Ambiguous categories (e.g., “misc_RNA”, “pseudogene”) and intermediate processing byproducts (e.g., miRNA loop regions) were excluded. Sequences shorter than 20 nt were also excluded. The final OpenRNA v1 dataset comprises 114,186,538 unique sequences, providing a non-redundant and biologically coherent training set.

#### 4.2.2 Data processing and tokenization

##### Sequence Clustering and Sampling

To reduce data redundancy and enhance training diversity, we clustered the 114M cleaned sequences using `MMseqs2` with parameters set to 50% sequence identity (`MIN_SEQ_ID=0.5`) and 80% coverage (`COVERAGE=0.8`). This clustering produced 17,350,557 clusters, with 56.19% being singleton clusters and 37.65% being ultra-small clusters (2–10 sequences), indicating substantial redundancy in the data.

To improve exposure of rare RNA types while maintaining data quality, we employed an inverse-sqrt weighted sampling strategy: the sampling probability *P_i_* for a cluster *i* is inversely proportional to the square root of its size 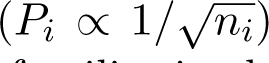, enabling the model to effectively learn complex sequence patterns from rare RNA families in the long-tail distribution.

##### Sequence Tokenization

Each sequence was annotated with two orthogonal labels: (i) RNA type, assigned using source database metadata to one of 15 RNA family categories; and (ii) taxonomic lineage, mapped to the seven-level Greengenes hierarchy (`d__domain; p__phylum; c__class; o__order; f__family; g__genus; s__species`) via NCBI Taxonomy lookups on the NCBI accession or organism field of each record. Sequences lacking a resolvable taxonomy at any rank were assigned partial lineage strings (with missing levels omitted), retaining the deepest available annotation. More than 90% of sequences received complete RNA-type and at least genus-level lineage annotations.

Nucleotide sequences were tokenized at single-nucleotide resolution using a character-level vocabulary consisting of four canonical RNA nucleotides (A, U, G, C), a 5 token denoting the 5^′^-to-3^′^ reading direction, a 3 token denoting the 3^′^-to-5^′^ reading direction (reverse complement), special tokens (`<bos<, <bos_glm>, <eos>, <eos_span>, <pad>, [MASK]`), 11 RNA-type conditioning tokens, and span sentinel tokens for GLM training. The tokenizer vocabulary has a total size of 114 tokens. Sequences exceeding 8,192 nt were truncated to fit EVA’s context window. Although OpenRNA v1 encompasses 15 RNA categories, only 11 are assigned dedicated conditioning tokens (mRNA, lncRNA, circRNA, snoRNA, snRNA, sRNA, tRNA, rRNA, piRNA, miRNA, and viral RNA); the remaining four categories (YRNA, scaRNA, ribozyme, and vaultRNA) lack sufficient representation to support reliable conditional generation and are therefore included in training without a category-specific conditioning token.

#### 4.2.3 Sequence diversity analysis within RNA families

To characterize intra-family sequence conservation across RNA types, we employed the Average Pairwise Identity (API) as the primary metric. Given a family of *N* sequences, API is defined as:

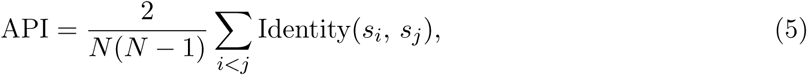

where Identity(*s_i_, s_j_*) is the fraction of identical positions between sequences *s_i_* and *s_j_* after global pairwise alignment, computed as the number of matched positions divided by total alignment length. API is a continuous measure in [0, 1] that captures subtle evolutionary trends missed by discrete clustering counts; unlike *k*-mer similarity or edit distance, it accounts for insertions and deletions through global alignment. High API indicates strong functional constraint; low API reflects greater sequence diversity or relaxed selective pressure.

Prior to API calculation, we applied 90% identity clustering to OpenRNA v1 using MM-seqs2 easy-linclust (`–min-seq-id 0.9, -c 0.8, –cov-mode 1`), reducing the 114M sequences to 37,447,972 representatives while preserving family diversity. API was then computed for 11 RNA families: mRNA, miRNA, lncRNA, circRNA, piRNA, sRNA, viral RNA, rRNA, tRNA, snRNA, and snoRNA.

Because the largest family (mRNA, *>*27M sequences) makes exhaustive *O*(*N* ^2^) pairwise computation infeasible, we adopted a non-parametric bootstrap resampling strategy: in each of *K* = 50 iterations, *n* = 500 sequences were drawn uniformly at random, aligned with MAFFT, and API was computed for the subsample. The final estimate is the mean over all iterations, with 95% confidence intervals and coefficients of variation reported as uncertainty measures. This reduces complexity from *O*(*N* ^2^) to *O*(*K · n*^2^). Sensitivity analysis confirmed that *n* = 500, *K* = 50 achieves a 95% CI width of approximately ±1% with coefficient of variation below 5%. API values for all families are reported in Supplementary Table S3 and Figure S2A.

#### 4.2.4 Phylogenetic tree construction of training data

To improve ecological coverage, multi-domain RNA representation, and lineage-level balance, a stratified sampling strategy was applied to characterize the ecological landscape of the dataset. Sequence redundancy was first reduced using MMseqs2 clustering at 50% sequence identity, and representative sequences were retained. To exclude biologically and computationally uninformative extremes, sequences outside the 80–4000 nucleotide length range were filtered to satisfy the model’s context length constraint. A hierarchical Hybrid Proportional-Minimum Sampling framework was then employed to preserve diversity across RNA functional categories, lineage-level ecological grouping, and sequence length distribution, enabling a multi-dimensional overview of dataset ecology. Ultimately, 1746 sequences were selected for visualization to illustrate the ecological and distributional landscape of the dataset. Phylogenetic overview trees were constructed sequentially using MAFFT for multiple sequence alignment, ClipKit for alignment trimming, and FastTree for tree inference.

#### 4.2.5 UMAP visualization of training data

To visualize the sequence-space diversity of OpenRNA v1, we computed 2-dimensional UMAP embeddings from species-level *k*-mer frequency profiles (*k* = 1–6) calculated over all RNA sequences. Prior to dimensionality reduction, feature vectors were re-weighted by domain (Archaea 5, Bacteria 2, Eukaryota 1) to compensate for imbalanced species counts, then *ℓ*_2_-normalized. UMAP was run with `n_neighbors= 30, min_dist= 0.8, spread= 1.5`, and cosine distance. Points are colored at the phylum level (Bacteria: green; Eukaryota: blue; Archaea: red), with color intensity reflecting the log-scaled number of sequences per species. The resulting projection is shown in Supplementary Figure S1B.

### 4.3 Model Benchmarking and Applications

Through self-supervised learning on RNA sequences alone, EVA learns universal representations that enable a wide range of downstream applications, from zero-shot fitness prediction to controllable generative design.

#### 4.3.1 Zero-shot Fitness Prediction

We benchmark EVA’s predictive capabilities across RNA, DNA, and protein modalities, evaluating whether the model’s likelihood scores correlate with experimental fitness.

##### Zero-shot ncRNA fitness prediction

We curated a benchmark of 13 deep mutational scanning (DMS) assays spanning three structured ncRNA classes—ribozymes, RNA aptamers, and transfer RNAs (tRNAs)—to evaluate EVA’s zero-shot fitness prediction capability (Supplementary Table S6). Assays were sourced from published DMS studies, covering diverse functional readouts including catalytic activity, fluorescence, and stability. For each assay, we computed the Spearman rank correlation coefficient (*ρ*) between per-variant EVA log-likelihood scores and the experimentally measured functional activities.

##### Zero-shot mRNA fitness prediction

We curated five datasets capturing diverse aspects of mRNA function, including protein-coding fitness and in-cell structural accessibility (Supplementary Table S7). Spearman rank correlation (*ρ*) between EVA log-likelihood scores and experimental measurements served as the primary evaluation metric, consistent with the ncRNA benchmark.

##### Zero-shot gene essentiality prediction

To evaluate whether EVA’s sequence likelihoods encode information about gene function at the organism level, we assessed its ability to predict gene essentiality in a zero-shot setting, following the methodology of Evo 2^19^.

For bacteria, we compiled a dataset of essential and non-essential genes from the Database of Essential Genes (DEG)^48^, cross-referenced with NCBI RefSeq genome annotations. Species were retained if they contained at least 10 essential and 10 non-essential annotated genes, yielding 42 bacterial species. For eukaryotes, we assembled essentiality annotations for 5 species (*Arabidopsis thaliana*, *Aspergillus fumigatus*, *Caenorhabditis elegans*, *Schizosaccharomyces pombe*, and *Saccharomyces cerevisiae*) from DEG^48^. The full list of species with their NCBI accession numbers is provided in Supplementary Table S9. Gene sequences were formatted with species-specific taxonomic lineage prefixes and RNA type tokens (e.g.,

`<bos<`|*lineage*;`<rna_mRNA>`|5*sequence* 3`<eos>`).

For each gene, we computed the autoregressive log-likelihood of the wild-type coding sequence under the model:

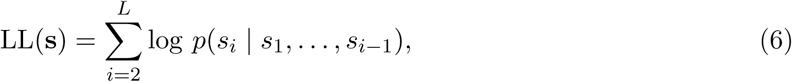

where **s** = (*s*_1_*, . . . , s_L_*) is the tokenized input sequence. We then constructed a mutant sequence by inserting five tandem premature stop codons (UAAUAAUAAUAAUAA; 15 nt) near the 5^′^ end of the coding region (at nucleotide position 12, or at one quarter of the gene length for short genes), and computed its log-likelihood LL(**s**^′^). The essentiality score was defined as the difference in log-likelihood between wild-type and mutant:

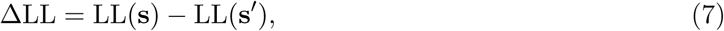

where a higher ΔLL indicates that the premature stop insertion is more disruptive to the model’s sequence likelihood, predicting greater essentiality. Sequences were truncated to a maximum of 8,192 tokens. We computed the area under the receiver operating characteristic curve (AUROC) for each species independently and averaged across species. We compared EVA (1.4B and 21M parameters) against Evo 2 (40B, 7B and 1B parameters) and two non-sequence baselines: for bacteria, a positional baseline based on distance from the chromosomal origin of replication (oriC); for eukaryotes, a GC content baseline.

##### Zero-shot protein fitness prediction

Although trained explicitly on RNA, EVA generalizes to protein fitness prediction. We evaluate this using 20 human protein DMS datasets that provide readily accessible nucleotide and protein sequences: ten from ProteinGym (ERBB2, NPC1, LYAM1, GLPA, PITX2, RBP1, RASH, CBPA2, PIN1, and DNJA1) and ten from DomainOme (Q9HC78, Q9UJQ4 [PF00096_383], Q8N1W1, Q9NU63, Q7Z5Q1, Q9ULZ3, Q13263, Q9UL15, O15151, and Q9UJQ4 [PF00096_627]). Full dataset identifiers and source databases are provided in Supplementary Table S8. We test whether the model captures protein-level evolutionary constraints via its exposure to coding sequences, computing Spearman rank correlation between per-variant EVA log-likelihood scores and experimentally measured fitness effects.

#### 4.3.2 Controllable Generative Design

Benefiting from our proposed training strategy and extensive data, EVA enables controllable rational design of diverse RNAs across all domains of life.

##### Conditional Autoregressive Generation

EVA supports fine-grained control over generation by conditioning on both RNA-type tags and taxonomic lineage tags. Conditioning on RNA type ensures the preservation of family-specific structural motifs (e.g., cloverleaf structures for tRNA). Further conditioning on taxonomic lineage enables species-aware design, such as optimizing mRNA codon usage to match the tRNA abundance of a specific host organism. This dual-conditioning capability is critical for practical applications like species-specific mRNA vaccine design. To assess how closely CLM-generated sequences resemble natural RNAs, we searched each generated sequence against the full OpenRNA v1 training corpus using MMseqs2 (nucleotide mode, `–search-type 3`, sensitivity *s* = 7.5, *k*-mer size 11, spaced *k*-mer mode, strand-aware, E-value ≤ 10, minimum alignment length 15 nt, up to 20 reported hits per query). For each generated sequence, we retained the database hit with the highest full-length weighted similarity, defined as:

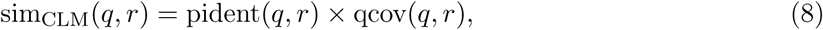

where pident(*q, r*) ∈ [0, 1] is the local percent sequence identity of the best-scoring alignment and qcov(*q, r*) ∈ [0, 1] is the fraction of the query sequence covered by that alignment. Multiplying identity by coverage penalises hits in which only a short, highly similar sub-region is matched, ensuring that sequences with poor full-length coverage receive low scores even when the aligned portion is highly conserved. If no database hit was found for a query, its similarity was set to 0. The resulting per-sequence similarity values were aggregated into the violin plots shown in Figure 3A.

##### Targeted Region Redesign (GLM)

Using the Generalized Language Modeling (GLM) capability, users can perform targeted redesign of specific RNA regions (e.g., IRES elements) while keeping flanking sequences fixed. This enables modular optimization of functional elements within larger RNA scaffolds. To quantify how accurately the model reconstructs masked sequence spans, we compared each model-generated infill against the corresponding ground-truth fragment using global pairwise alignment. Alignments were performed with Biopython’s `PairwiseAligner` in global mode (match score +1, mismatch score 1, gap-open penalty 2, gap-extend penalty 0.5), implementing the Needleman–Wunsch algorithm. The reconstruction similarity for a generated fragment *ŝ* relative to its ground truth *s*^∗^ was computed as:

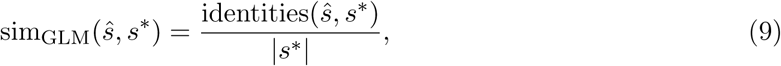

where identities(*ŝ, s*^∗^) is the number of identical nucleotide pairs in the optimal global alignment and *s*^∗^ is the length of the ground-truth fragment. Normalising by ground-truth length rather than alignment length ensures that insertions in the generated sequence are penalised. Per-span similarity values were aggregated into the violin plots shown in Figure 3B, stratified by RNA class.

##### Secondary structure statistics for distributional comparison

To compare the biophysical properties of generated sequences against natural RNAs at scale, secondary structures were predicted using RNAfold from the ViennaRNA package (default parameters). RNAfold was chosen over slower folding tools because the distributional comparisons require predicting structures for large numbers of sequences (up to 200 generated sequences and thousands of natural reference sequences per RNA class or species), making computational efficiency essential. For each sequence, RNAfold provides the minimum free energy (MFE) dot-bracket secondary structure and the corresponding MFE value. The following structural statistics were extracted from the predicted dot-bracket string: MFE (kcal/mol), reflecting overall thermodynamic stability; stem/helix fraction, the proportion of nucleotides in Watson–Crick base pairs; loop fraction, the proportion of unpaired nucleotides; and base-pairing rate, the ratio of paired to total nucleotides summarizing global structural compactness. Distributions of these statistics for generated sequences (*n* = 200 per condition) and natural sequences were compared using kernel density estimates and visualized as overlaid density plots, enabling direct assessment of whether EVA’s generative distribution recapitulates the structural properties of natural RNAs.

#### 4.3.3 Zero-shot tRNA Generation and Structural Evaluation

To evaluate EVA’s ability to capture three-dimensional structural constraints from sequence alone, we performed zero-shot conditional generation of tRNAs by conditioning the model on the tRNA type tag without any structure-based supervision.

For each generated sequence, we predicted its three-dimensional structure using AlphaFold3 and compared it against a curated set of 32 experimentally resolved monomeric tRNA structures retrieved from the RCSB Protein Data Bank (PDB IDs: 1EHZ, 1EVV, 1FIR, 1I9V, 1QZB, 1TN1, 1TN2, 1TRA, 1VTQ, 1YFG, 2K4C, 2TRA, 3A3A, 3BBV, 3CW5, 3CW6, 3L0U, 3TRA, 4P5J, 4TNA, 4TRA, 5L4O, 6CU1, 6TNA, 6UGG, 6Y3G, 7EQJ, 7KJU, 7VNW, 8UPT, 8UPY, 8V1H). Two metrics were computed for each generated sequence. For structural similarity, the AlphaFold3-predicted structure was compared against the same curated set of 32 RCSB tRNA structures and the maximum TM-score was taken. For sequence novelty, we randomly sampled 10,000 tRNA sequences from OpenRNA v1 and computed pairwise sequence identity via Needleman–Wunsch global alignment; the maximum identity across all 10,000 comparisons was taken as the sequence similarity score, with lower values indicating greater novelty relative to the training data. Representative generated sequences were additionally superimposed onto their closest RCSB structural matches for visual inspection of tertiary fold recapitulation.

#### 4.3.4 Fine-tuning for Specialized RNA Types

To extend EVA’s applicability to RNA families underrepresented in the training data, we implemented a lightweight fine-tuning strategy that preserves pre-trained RNA knowledge while adapting the generative distribution to user-defined functional RNA classes. We introduced a dedicated `Y_RNA` conditioning token reserved for user-defined RNA types; during fine-tuning, all sequences in the specialized dataset are assigned this token, enabling the model to associate a distinct generative mode with the target class without interfering with the 11 pre-trained RNA-type distributions. Two case studies were pursued: (i) RNA aptamers, synthetically engineered fluorogenic molecules absent from natural sequence databases, fine-tuned on *n* = 30 high-fluorescence sequences; and (ii) IscB omegaRNAs, CRISPR guide RNAs from a metagenomic Cas-related system, fine-tuned separately on high-editing-efficiency sequences from two guide variants (m16: *n* = 42; m17: *n* = 44). Starting from the pre-trained EVA checkpoint, we performed continued autoregressive language-model training on each specialized dataset using a reduced learning rate and a small number of epochs, sufficient to shift the generative distribution while retaining general RNA sequence knowledge.

After fine-tuning, sequences were sampled autoregressively conditioned on the Y_RNA token. For omegaRNA design, two independent models (one per guide variant) were each used to generate large libraries (m16: *n* = 1,800; m17: *n* = 1,595), and generated sequence lengths were verified against the expected length range of natural IscB omegaRNAs. To characterize the edit distance between generated sequences and wildtype, we counted the number of mutated positions in each generated aptamer sequence (*n* = 93) relative to the wildtype and compared the resulting distribution to that of variants catalogued by deep mutational scanning (DMS), establishing whether fine-tuned generative models produce sequences with biologically plausible divergence levels.

Representative designed sequences from each case study were submitted to AlphaFold3 for complex structure prediction. For *ω*RNA designs, ternary complexes comprising the designed guide RNA, IscB protein, and target DNA substrate were predicted using AlphaFold3 in MSA mode with the experimentally resolved IscB–*ω*RNA–DNA structure (RCSB PDB: 7UTN) as a structural template. To move beyond global confidence scores, we derived five domain-level geometric criteria from the IscB structural literature^25,26^ and applied them as pass/fail filters across all predicted complexes (Supplementary Table S13). The five criteria cover: (1) HNH domain distance to the guide RNA, verifying productive nuclease positioning for target-strand cleavage; (2) contact of all four PLMP residues with the *ω*RNA (heavy-atom distance *<* 7 Å), required for complex stabilization; (3) straightness of the bridge helix (BH) *α*-helix; (4) fraction of the BH surface wrapped by the *ω*RNA, reflecting scaffold integrity; and (5) WED domain distance to the PAM-proximal DNA phosphate backbone, confirming correct PAM engagement. Designs satisfying all thresholds were shortlisted as structurally viable candidates and visualized from two opposing viewpoints (0° and 180° rotation) to confirm overall fold preservation.

### 4.4 Design and Optimization of RNA Vaccines

We applied EVA to optimize two major classes of RNA vaccines: linear mRNA vaccines and circular RNA (circRNA) vaccines.

#### 4.4.1 Species-aware mRNA Codon Optimization

We formulated codon optimization as a synonymous substitution search over CDS sequences, jointly optimizing host compatibility and thermodynamic stability. Rather than relying on the Codon Adaptation Index (CAI), which treats each codon position independently, we use EVA’s species-conditioned log-likelihood as the primary scoring signal. Prepending the target species tag causes EVA to score each sequence against the host-specific evolutionary manifold, capturing global context including UTR compatibility, structural constraints, and organism-specific preferences in a single data-driven objective.

##### Optimization Algorithm

Optimization proceeds via sliding-window beam search combined with simulated annealing over synonymous coding sequences, starting from *P* = 8 candidate CDS sequences. At each iteration, *B* = 8 sequences are drawn as modification targets. For each, a window of *W* = 8 codons is placed at a random CDS position, and EVA performs beam search (width *k* = 10) over valid synonymous assignments given the upstream context:

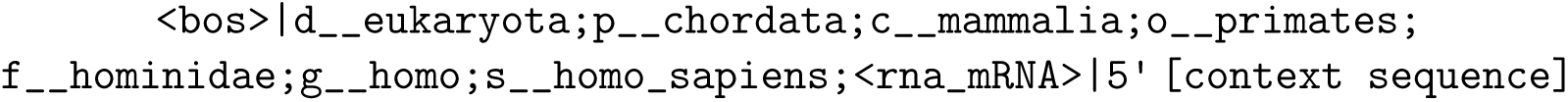

where `<rna_mRNA>` is replaced by `<rna_circRNA>` for circRNA ORF optimization. The top-*k* beam candidates are scored by LinearFold MFE; the lowest-MFE candidate is accepted via the Metropolis criterion (Δ*E <* 0 unconditionally; otherwise with probability exp(−Δ*E/T*)). Temperature decays exponentially from *T*_init_ = 2.0 to *T*_min_ = 0.01 (*γ* = 0.995) over *N* = 400 iterations. As an ablation, a *Random (MFE-only)* baseline replaces EVA beam search with uniform random synonymous sampling while retaining the same Metropolis criterion.

##### Evaluation

Optimized sequences were evaluated on CAI (post hoc, from human codon usage frequencies as the geometric mean of per-codon RSCU weights) and MFE (ViennaRNA; lower = more stable). We benchmarked on five vaccine candidates: SARS-CoV-2 Spike, Influenza PR8 HA (A/PR/8/34 H1N1), VZV gE, RABV-G, and HIV envelope protein.

#### 4.4.2 ORF Optimization and IRES Generation for circRNA Vaccines

For circRNA vaccines, which lack 5^′^ caps and poly(A) tails, design involves optimizing the ORF and engineering the Internal Ribosome Entry Site (IRES) for cap-independent translation.

##### ORF Optimization

The circRNA ORF was optimized using the same beam search–simulated annealing pipeline as for linear mRNA. We evaluated four systems: SARS-CoV-2 (Group-I intron and T4 ligase cyclization), Rabies, and CAR-T circRNAs.

##### De Novo IRES Redesign via GLM

We used EVA’s GLM capability to redesign IRES elements in a fill-in-the-middle fashion. The original CVB3 IRES is treated as a masked span, with flanking ORF and permuted-intron-exon (PIE) sequences held fixed as left and right context. A `<span_0>` sentinel is placed at the IRES position, conditioned on the circRNA type and *Homo sapiens* lineage tags; EVA then samples IRES candidates autoregressively from both flanking contexts. Candidates are evaluated on two key structure-based metrics (ViennaRNA/LinearFold): ribosome accessibility (unpaired probability of the AUG region) and inhibitory long-range interaction fraction (IRES base pairs with distal ORF or PIE sequences), and benchmarked against the original CVB3 IRES.

### 4.5 Model Interpretability with Sparse Autoencoders

#### 4.5.1 Sparse Autoencoder Training

To decompose dense, distributed representations into interpretable features, we trained sparse autoencoders (SAEs) on model internal activations, following recent advances in mechanistic interpretability^37^. We trained SAEs on two EVA model scales (145M and 1.4B parameters), each with an architecture matched to its downstream analysis objective: a ReLU SAE with soft *L*_1_ sparsity on EVA 145M for global feature-space visualization and domain-level segregation analysis, and a BatchTopK SAE with hard sparsity on EVA 1.4B for per-nucleotide functional element annotation in therapeutic RNA constructs. Both were trained on layer 13 activations extracted from one million RNA sequences sampled from a curated multi-species corpus. Layer 13 was empirically selected as an intermediate layer capturing the transition from local sequence features to higher-order biological abstractions.

##### ReLU SAE with *L*_1_ sparsity

For the 145M-parameter model (hidden dimension *d* = 448), we trained a ReLU-based SAE following the InterPLM framework^49^. The encoder maps the centered input to an overcomplete latent space of dimensionality *D* = 3,584 (8× expansion):

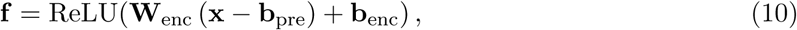

and the decoder reconstructs the input as **x**^ = **W**_dec_ **f** + **b**_pre_. The training loss is:

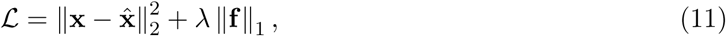

with *λ* = 0.06 linearly warmed up over the first 5% of training steps. The model was optimized with ConstrainedAdam^49^ (learning rate 2 × 10^−4^, batch size 16,384 tokens, 150,000 steps) with periodic dead-neuron resampling. This SAE is used for global feature-space visualization (Section 4.5.3), where continuous activation densities across large sequence corpora are required.

##### BatchTopK SAE

For the 1.4B-parameter model (hidden dimension *d* = 1,024), we adopted a BatchTopK SAE architecture ^50^ enforcing hard sparsity constraints. The encoder maps the centered input to *D* = 8,192 latent dimensions (8× expansion), with a BatchTopK operation fixing exactly *k* = 32 active features per token on average. The training loss combines reconstruction error with an auxiliary dead-latent penalty^50^ (*α* = 0.03) to encourage full utilization of latent capacity. Optimization used Adam with a trapezoidal learning rate schedule (peak 1 × 10^−4^, gradient norm clipped at 1.0). The final SAE achieved reconstruction loss ~ 0.76 and 100% feature utilization across all 8,192 dimensions. This SAE is used for functional element annotation of therapeutic RNA constructs (Section 4.5.2), where a fixed per-token sparsity level (*L*_0_ = 32) ensures comparability of activation magnitudes across nucleotide positions.

#### 4.5.2 Functional Element Annotation of Therapeutic RNA Constructs

To assess whether SAE features capture functionally meaningful regulatory and coding elements in engineered therapeutic constructs, we applied the BatchTopK SAE to annotated circRNA and mRNA vaccine sequences. For each construct, we passed the full sequence through EVA 1.4B, extracted layer 13 activations, and projected them through the SAE encoder (*k* = 32), yielding a per-nucleotide activation matrix **A** ∈ ℝ*^L^*^×^*^D^*.

##### Region-specific feature selection

For each annotated functional region *r* (start *s_r_*, end *e_r_*), we identified the SAE feature best discriminating the target region from the remainder of the sequence via a penalty-weighted score:

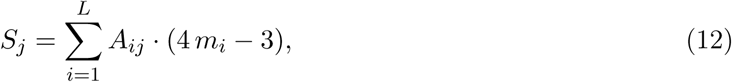

where *m_i_*= 1 for *s_r ≤_ i < e_r_* and 0 otherwise. This assigns weight +4 to in-region activations and 3 to off-target activations. Features were selected by maximizing coverage (fraction of in-region positions where the feature is active) subject to a leakage threshold of *<* 50% (ratio of out-of-region to in-region activation); the highest-scoring feature was used when no feature met this criterion. Functional regions shorter than 50 nt were excluded.

##### circRNA constructs

We analyzed two circRNA vaccine candidates: a RABV-G construct (2,348 nt, containing a CVB3 IRES and RABV-G ORF) and a SARS-CoV-2 RBD construct (1,695 nt, containing a CVB3 IRES and SP+RBD coding sequence) ^27^. Each construct was visualized as a polar plot reflecting its covalently closed topology, with per-nucleotide activation magnitudes shown as radial bars.

##### mRNA constructs

We analyzed four codon-optimized mRNA constructs sharing identical UTRs (5^′^UTR: 64 nt; 3^′^UTR: 94 nt) encoding distinct viral antigens: HIV-1 gp160 (2,571 nt CDS), influenza A/PR8 hemagglutinin (1,701 nt CDS), RABV-G (1,575 nt CDS), and varicella-zoster virus glycoprotein E (1,749 nt CDS). Applying the penalty-weighted scoring independently to each construct, the same three features were selected across all four mRNAs: feature f/1998 for the 5^′^UTR, feature f/4512 for the CDS, and feature f/6236 for the 3^′^UTR. This consistency across constructs with divergent coding sequences provides evidence that these features encode generalizable functional concepts rather than sequence-specific motifs.

#### 4.5.3 UMAP Visualization of SAE Feature Space

To characterize the global organization of learned features and assess domain-level segregation, we performed UMAP visualization of the SAE decoder geometry, following the approach of Evo 2^19^. Each SAE feature *j* is associated with a decoder direction d*_j_* ∊ ℝ*^d^* (the *j*-th column of **W**_dec_). All *D* = 3,584 decoder directions from the 145M ReLU SAE were normalized to unit *ℓ*_2_-norm, and UMAP was applied to the resulting *D×d* matrix (cosine distance, *n*_neighbors_ = 15, min_dist = 0.1).

To assign domain-level biological interpretation, we computed differential activation frequencies between eukaryotic and prokaryotic sequences. We sampled 100,000 sequences from each domain from the training corpus, passed them through EVA and the SAE at layer 13, and recorded the fraction of tokens at which each feature was active (activation *>* 0), yielding per-feature densities 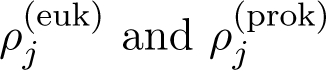. The differential score is:

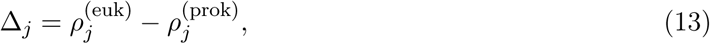

where Δ*_j_ >* 0 indicates a eukaryote-biased feature and Δ*_j_ <* 0 indicates a prokaryote-biased feature.

#### 4.5.4 Mixture-of-Experts Routing Analysis across RNA Biotypes

To assess whether different RNA biotypes induce systematically different expert routing patterns, we analyzed the routing behavior of EVA (145M parameters; 16 transformer layers, 8 experts per MoE layer, top-2 gating) using up to 50,000 sequences per RNA biotype sampled from a curated multi-species corpus spanning eight classes: mRNA, rRNA, tRNA, lncRNA, circRNA, miRNA, snoRNA, and snRNA. Sequences were truncated to 2,048 tokens and processed in mini-batches of 16.

##### Router probability extraction

For each MoE layer, we registered a forward hook on the gating module to capture softmax-normalized routing probabilities before expert dispatch. Probabilities were averaged across all tokens of a given RNA biotype to obtain a per-biotype, per-layer routing profile **p̂** ∈ ℝ*^E^*.

##### Routing specialization quantification

To quantify RNA-type-specific expert specialization at each layer, we computed the mean standard deviation of routing probabilities across RNA biotypes:

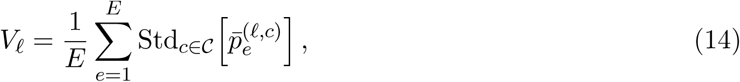

where 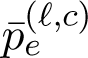 is the mean routing probability for expert *e* at layer *ℓ* over all tokens of RNA biotype *c*, and denotes the set of eight RNA classes. Higher *V_ℓ_* values indicate greater divergence in expert utilization across RNA types. The analysis was performed both without lineage prefix (raw sequence only) and with species-specific taxonomic prefix prepended, to assess how lineage conditioning influences expert routing.

## Acknowledgements

This study was supported by the National Natural Science Foundation of China (62041209), the Natural Science Foundation of Shanghai (24ZR1440600), the Science and Technology Commission of Shanghai Municipality (24510714300), and the Lingang Lab Fund (LGL-8888). We thank Shuya Li, Xiaoliang Shi, Bibi Zhang and Xiaoyu Chen for helpful discussions. We thank the Lingang Laboratory for providing computational resources. We thank Chaoxiang Lan for valuable advice on model training. We thank Lan Li for her aesthetic guidance and suggestions on the schematic figures. We thank Wenhao Yang, Runze Ma, and Zhongmin Li for their support on benchmarking. We thank Aiyuan Zhangxv, Hanlin Yu, Xiangwen Gao, Xinyi Zhang and Songan Zhang for their moral support.

## Author Contributions

S.Z. conceived the project. S.Z. and Y.H. supervised the project. G.L. and Y.H. designed the model architecture. Y.H. developed the first version of the training code; G.L. contributed to the training code. Y.H. developed the first version of the inference code; G.L., W.X., X.M., and Yij.H. contributed to the inference code. G.L., A.C., and Y.H. curated and processed the pretraining dataset. Y.H. conducted model pretraining, mid-training, needle-in-a-haystack evaluation, and related evaluations. Y.H., X.M., Yij.H., A.C., G.L. and Y.T. performed zero-shot mutational effect prediction. W.X. and Y.H. evaluated the model’s generative capabilities and performed the associated visualizations. Y.H. wrote the fine-tuning code and performed model fine-tuning. Y.H. and Y.X. conducted tRNA and RNA aptamer design, analysis, and visualization. Y.H., L.Z., and Y.X. performed CRISPR (IscB omegaRNA) data collection, design, analysis, and visualization. Y.H. organized the mRNA and circRNA vaccine datasets. W.X. wrote the codon optimization code for mRNA and circRNA vaccines, conducted codon optimization experiments, and performed the associated visualizations. Y.H. wrote the circRNA IRES generation code and conducted circRNA IRES design, analysis, and visualization. A.C. and Y.H. implemented and trained the sparse autoencoders (SAEs); A.C. performed SAE feature discovery and visualization. Y.C., Z.W., and Y.H. developed the model introduction and usage website; G.L. provided the backend model interface and code. Q.S. performed evolutionary analysis and visualization of the training data. M.C. and Y.H. conceived and created all conceptual figures. Y.H. wrote the first draft of the manuscript. All authors contributed to writing the final manuscript.

## Data and Materials Availability

To foster reproducibility and accelerate advancement in the RNA modeling community, we make all resources associated with this work publicly available.

- The OpenRNA dataset used to train EVA is available at: https://huggingface.co/datasets/GENTEL-Lab/OpenRNA-v1-114M
- All EVA model checkpoints spanning four scales (21M, 145M, 437M, and 1.4B parameters) are openly accessible at: https://huggingface.co/GENTEL-Lab/EVA
- Comprehensive training pipelines for both mixture-of-experts (MoE) and dense architectures, including pretraining, midtraining, and evaluation protocols, can be found at: https://github.com/GENTEL-lab/EVA/tree/main/training
- Fine-tuning workflows for downstream RNA design tasks are available at: https://github.com/GENTEL-lab/EVA/tree/main/finetune
- For interpretability analysis, we release Sparse Autoencoder (SAE) features and associated notebooks enabling mechanistic insights into EVA’s internal representations: https://github.com/GENTEL-lab/EVA/tree/main/notebooks/interpretability_analysis
- Practical tools for RNA engineering applications, including fitness prediction, causal language model (CLM)/generalized language model (GLM) design, and directed evolution workflows, are provided at: https://github.com/GENTEL-lab/EVA/tree/main/tools
- Complete benchmark datasets and experimental results for RNA generation and codon optimization are archived at: https://zenodo.org/records/19027142

## Supplementary Materials

- Supplementary Figures S1 to S10.
- Supplementary Tables S1 to S16.
- Supplementary Data.

OpenRNA v1 dataset: https://huggingface.co/datasets/GENTEL-Lab/OpenRNA-v1-114M

