## Supplemental figures for "A Long-Context Generative Foundation Model Deciphers RNA Design Principles"

### Supplementary Figures

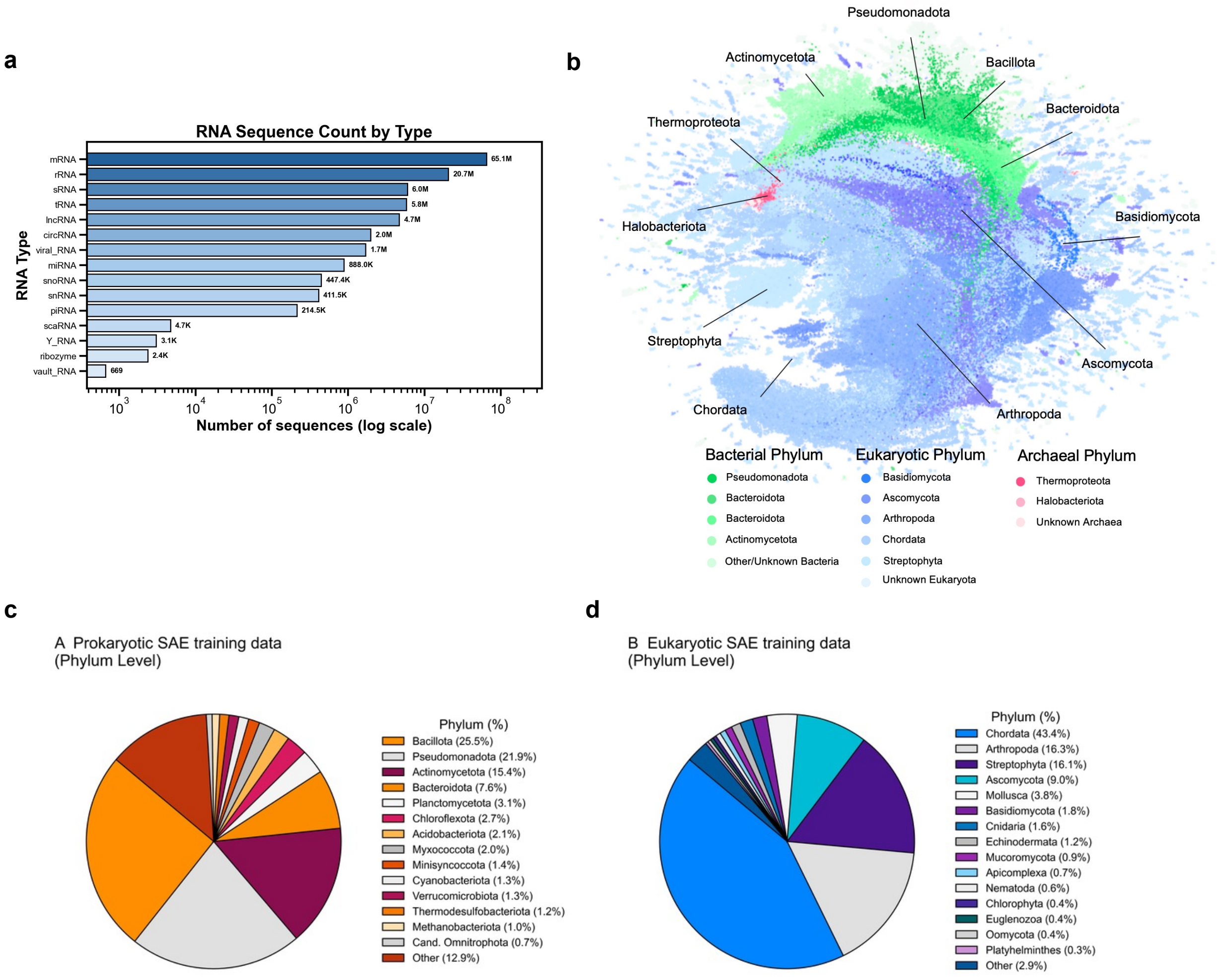


Figure S1: **Distribution of training data across RNA types and taxonomic groups.** (a) Proportion of different RNA types in the training dataset, showing the abundance distribution across 15 RNA categories. (b) Species composition in the training dataset visualized by UMAP, where each point represents a single RNA sequence, revealing the taxonomic diversity spanning all domains of life. (c) Phylum-level distribution of training data for prokaryotic (left) and eukaryotic (right) sequences, demonstrating comprehensive coverage across major taxonomic lineages.

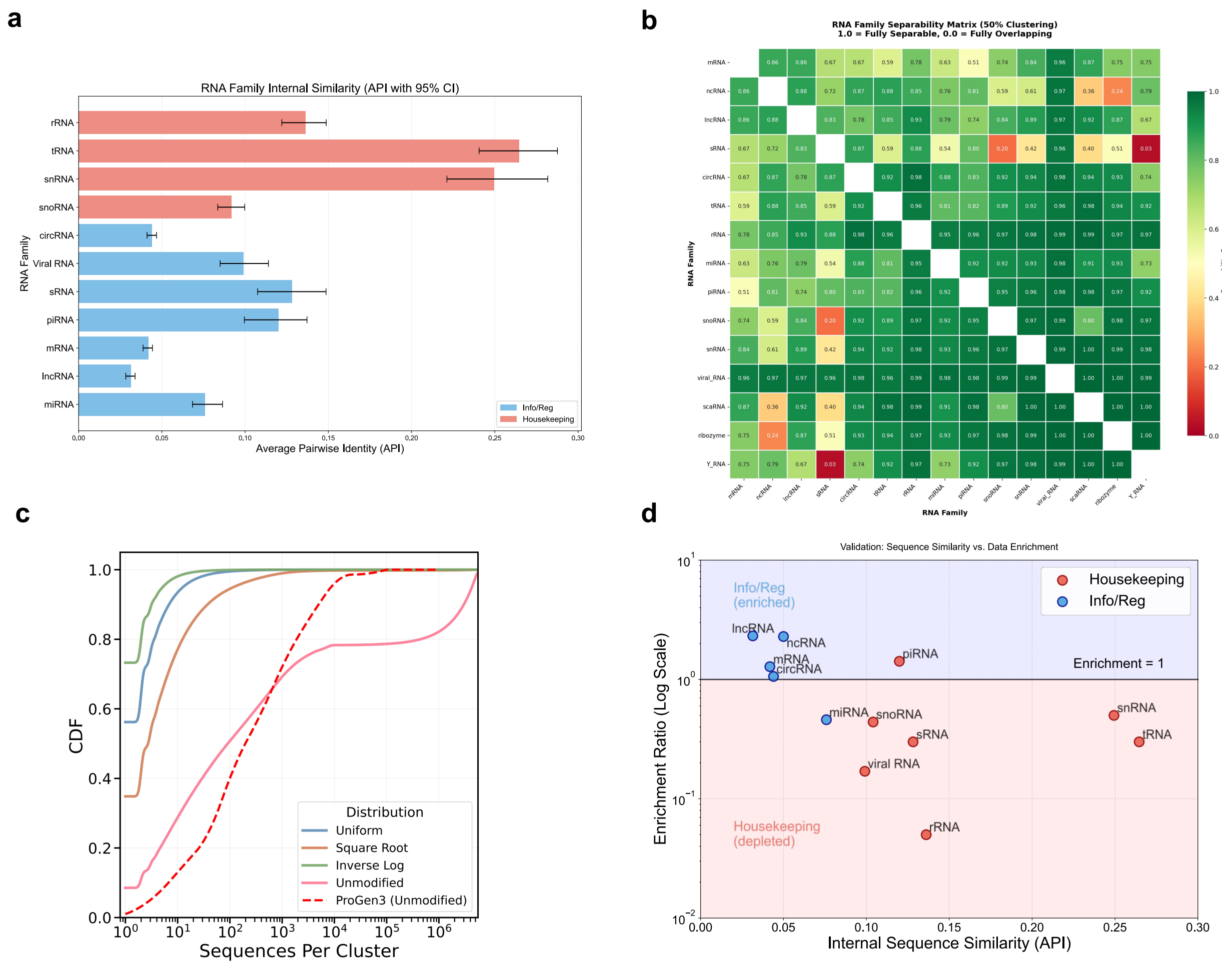

Figure S2: **Sequence conservation analysis and evolutionary conservation-based sampling strategy.** (a) Intra-RNA-type sequence similarity across different RNA categories. Housekeeping RNAs (e.g., rRNA, tRNA) exhibit high internal sequence conservation, while information-encoding and regulatory (reg) RNAs show lower conservation, indicating greater sequence diversity that requires more model attention during training. (b) Inter-RNA-type sequence similarity matrix showing substantial divergence between different RNA classes, highlighting the necessity of RNA-type conditioning tokens to enable efficient learning across diverse RNA families. (c) Cumulative distribution function (CDF) of cluster sizes under different sampling strategies. RNA sequences exhibit more severe data imbalance compared to proteins, characterized by both numerous extremely large clusters and many small clusters (potentially due to lower evolutionary conservation in RNA). Without sampling, direct training leads to overfitting on abundant clusters. Different sampling methods represent trade-offs between incorporating small clusters (potential noise) and down-weighting extremely large clusters. Square-root weighting achieves the best balance and performance. (d) Changes in RNA-type abundances after evolutionary conservation-based sampling. Housekeeping RNAs are down-sampled, allowing the model to focus more on learning complex sequence patterns in information-encoding (info) and regulatory (reg) RNA classes.

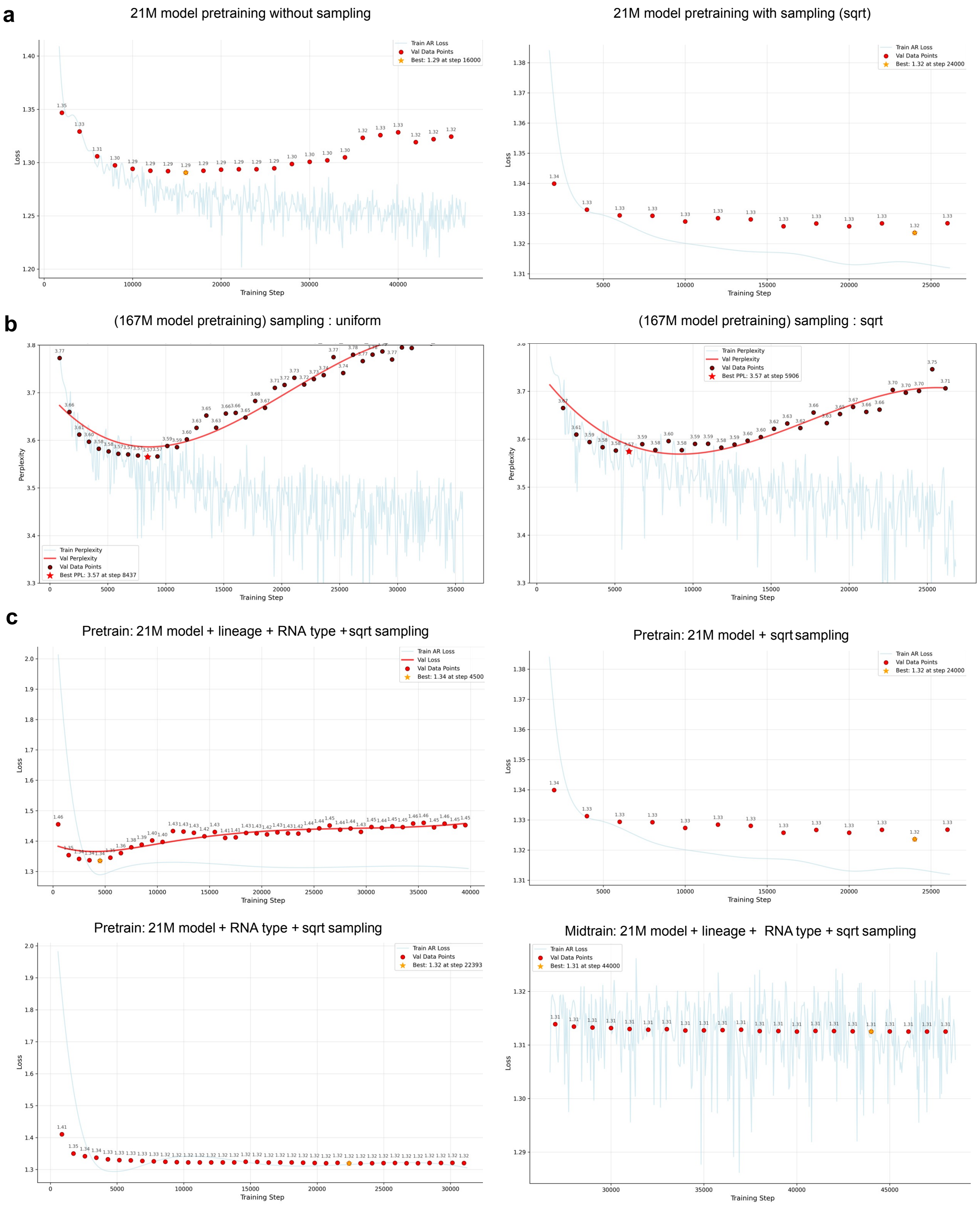


Figure S3: **Impact of sampling strategies and lineage conditioning timing on model training.** (a) Comparison of training dynamics with and without evolutionary conservation-based sampling, demonstrating that this strategy effectively prevents overfitting to abundant clusters. (b) Evaluation of different sampling strategies (uniform vs. square-root weighting). Square-root weighting achieves better performance by preventing overfitting to extremely large clusters while maintaining reasonable coverage of diverse RNA families. (c) Effect of lineage (taxonomic) conditioning timing on model performance. Introducing species-level information during pre-training leads to overfitting. Only when the model first learns universal RNA grammar during the pre-training stage, and then incorporates species-specific constraints in the mid-training stage, can overfitting be effectively prevented and optimal performance achieved.

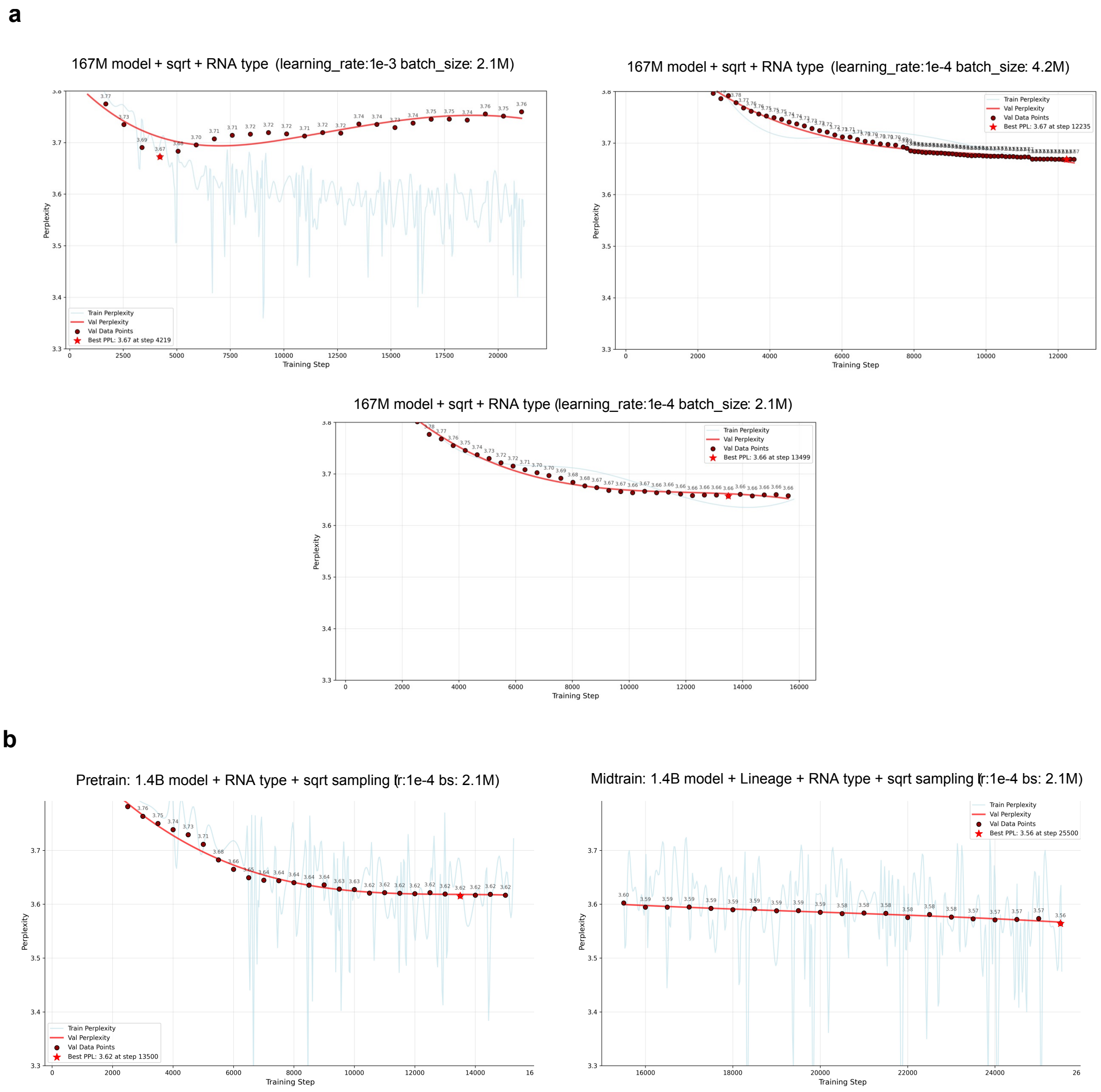


Figure S4: **Hyperparameter optimization and impact of mid-training lineage conditioning on perplexity.** (a) Learning rate exploration with fixed batch size. RNA foundation models may require smaller learning rates compared to DNA and protein foundation models with similar Transformer MoE architectures. For reference, ProGen3 (Transformer MoE, batch size ∼2M) uses a learning rate of $5 \times 10^{-4}$, and Evo2 (batch size 2.1M) uses $3 \times 10^{-4}$. Our experiments demonstrate that the optimal learning rate for RNA is $1 \times 10^{-4}$, suggesting distinct optimization dynamics for RNA sequence modeling. (b) Effect of incorporating species-level information during mid-training. The introduction of lineage (taxonomic) conditioning in the mid-training stage further refines the RNA manifold by disentangling species-specific constraints from universal RNA grammar, resulting in reduced perplexity and improved model performance.

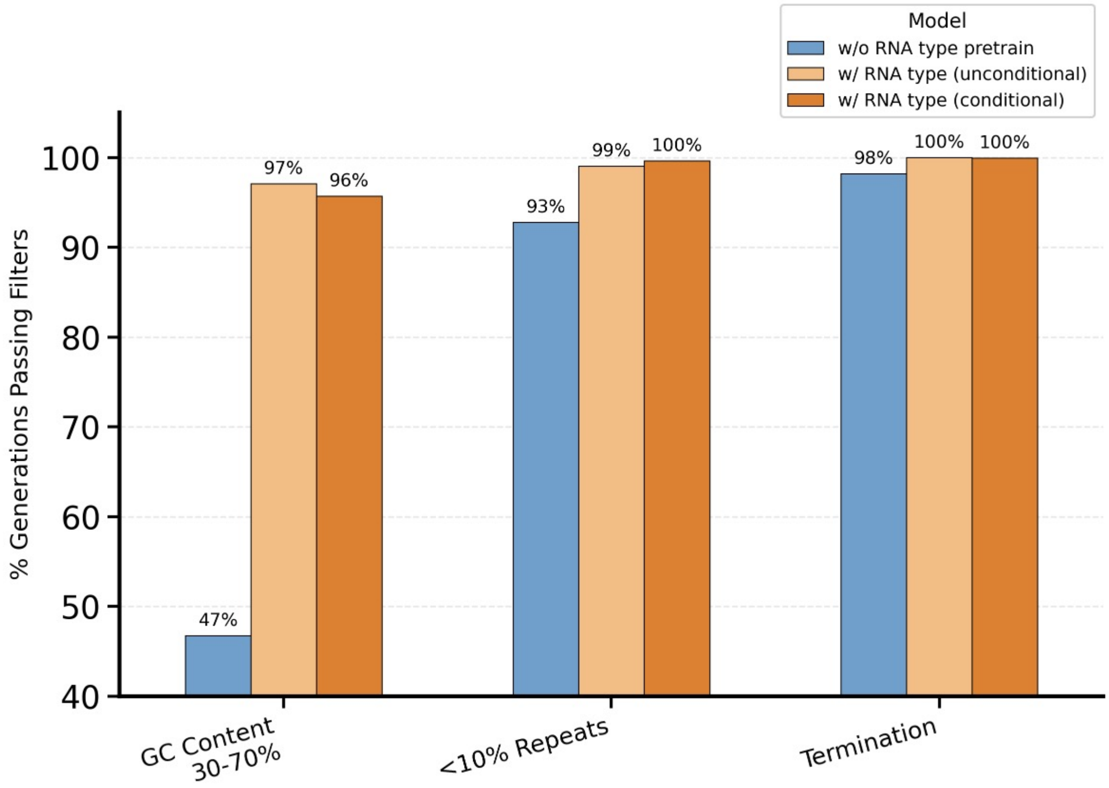


Figure S5: **Effect of RNA-type token conditioning on generation quality during pre-training.** Incorporating RNA-type tokens during pre-training substantially improves the quality of generated sequences, as measured by three criteria: (1) **GC content validity**: the sequence GC content falls within the biologically plausible range of 30%–70%; (2) **Semantic completeness**: the model generates a valid end-of-sequence (`<eos>`) token, indicating a properly terminated sequence; (3) **Low repetitiveness**: repetitive fragments constitute no more than 10% of the sequence. Repetitive generation is a common failure mode in biological language models. Repetitive fragments are defined as unit motifs of length 1–6 bp repeated at least 4 consecutive times, covering 5,440 detectable combinations in total (see Table S10 for details). As shown in the figure, pre-training without RNA-type token conditioning (blue bars) leads to a notably higher proportion of sequences with invalid GC content and excessive repetitive fragments, highlighting the critical role of RNA-type tokens in guiding the model toward biologically plausible sequence generation.

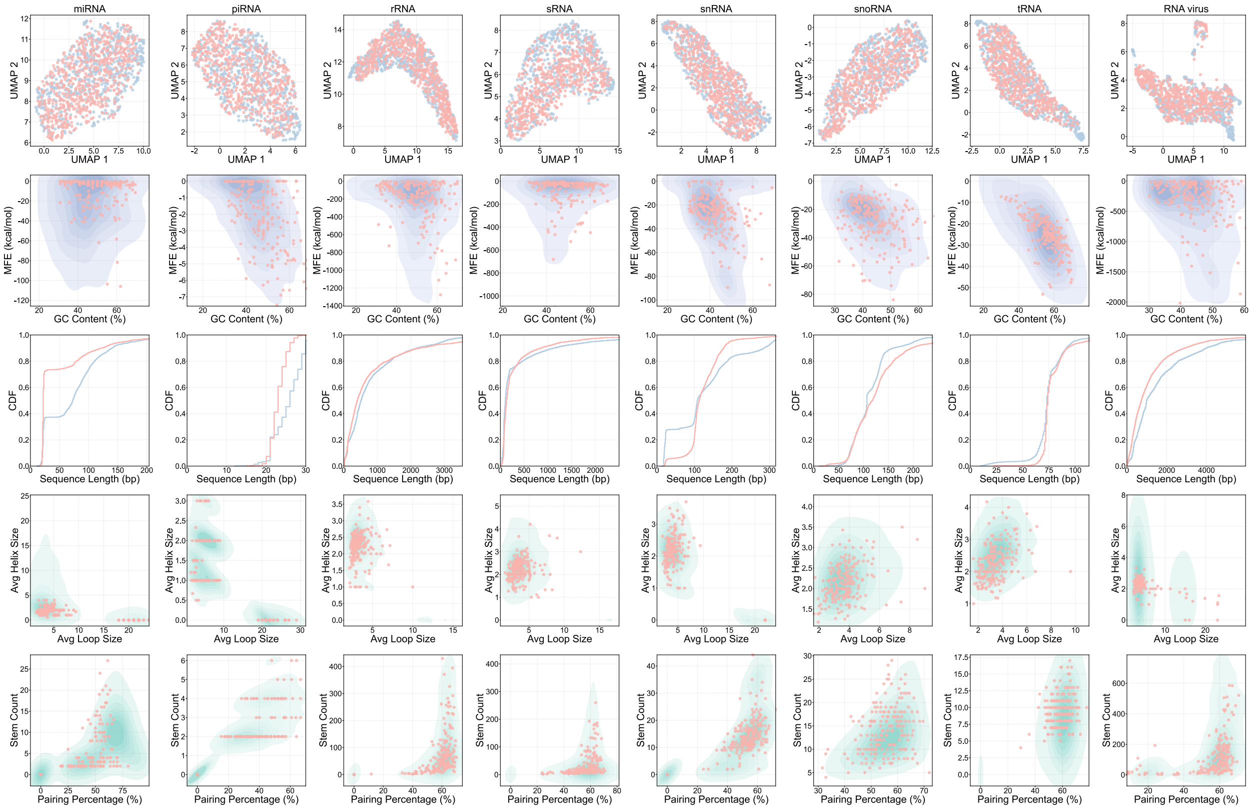


Figure S6: **RNA-type conditioned generation across all remaining RNA classes.** Distribution comparisons between EVA-generated sequences ($n = 200$, red dots) and natural RNAs (blue and green contours) for the 11 RNA classes not shown in the main text, evaluated across sequence-level metrics (UMAP, GC content) and structural metrics (MFE, secondary-structure statistics). Generated sequences closely track the natural sequence manifold across all RNA types, demonstrating that EVA accurately learns the distributional properties of each RNA class at both the sequence and structural levels.

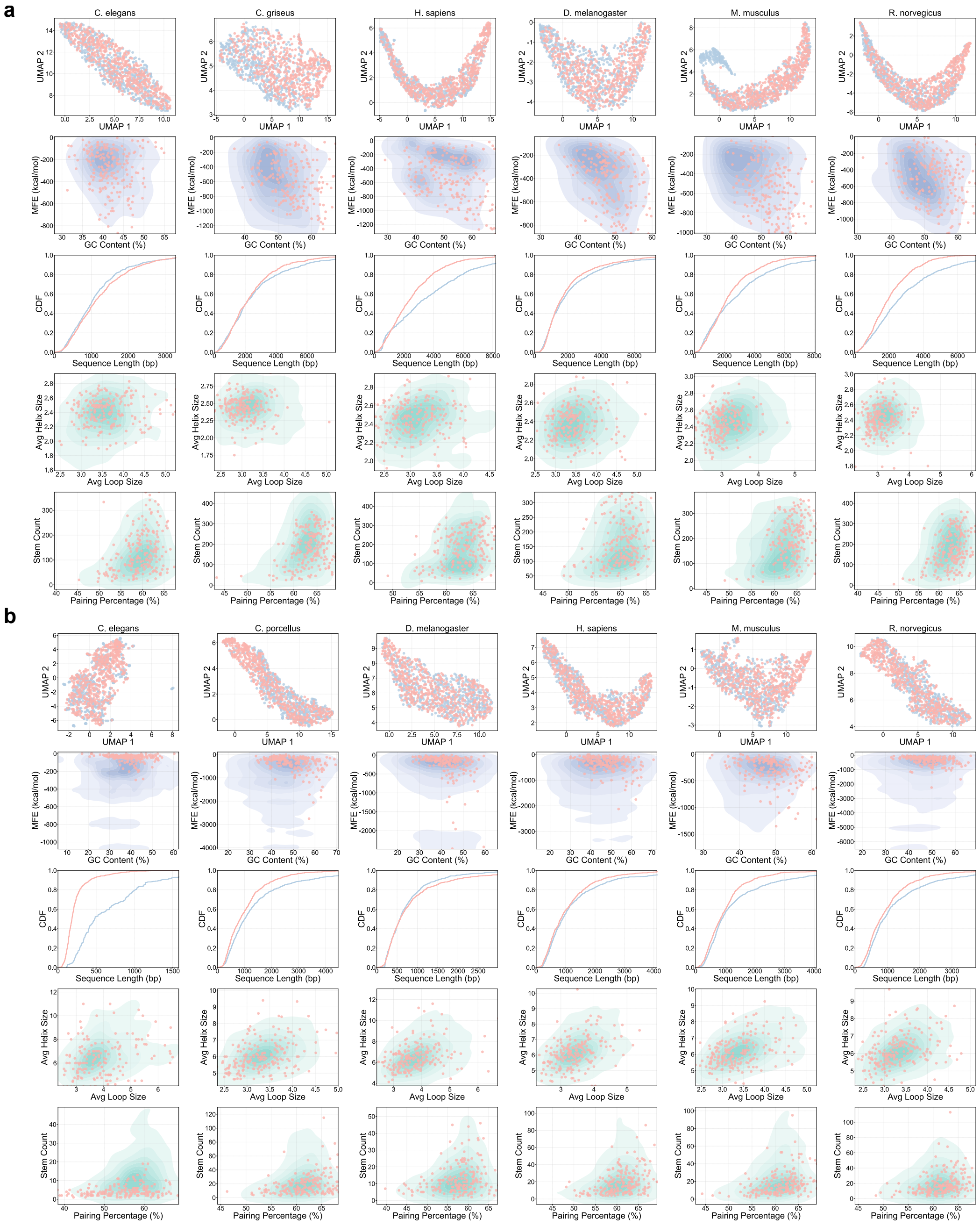

Figure S7: **Species-conditioned generation for mRNA and lncRNA across six species.** Distribution comparisons between EVA-generated sequences ($n = 200$, red dots) and natural RNAs (blue and green contours) conditioned on six representative species (*C. elegans*, *C. griseus*, *H. sapiens*, *D. melanogaster*, *M. musculus*, and *R. norvegicus*), evaluated across sequence-level metrics (UMAP, GC content) and structural metrics (MFE, secondary-structure statistics). (a) Species-conditioned generation within the mRNA class. Generated sequences closely recapitulate the species-specific sequence and structural distributions of natural mRNAs. (b) Species-conditioned generation within the lncRNA class. EVA similarly captures the species-specific manifold of lncRNAs across all six organisms, demonstrating that the model generalizes its controllable generation capability beyond mRNA to other RNA classes.

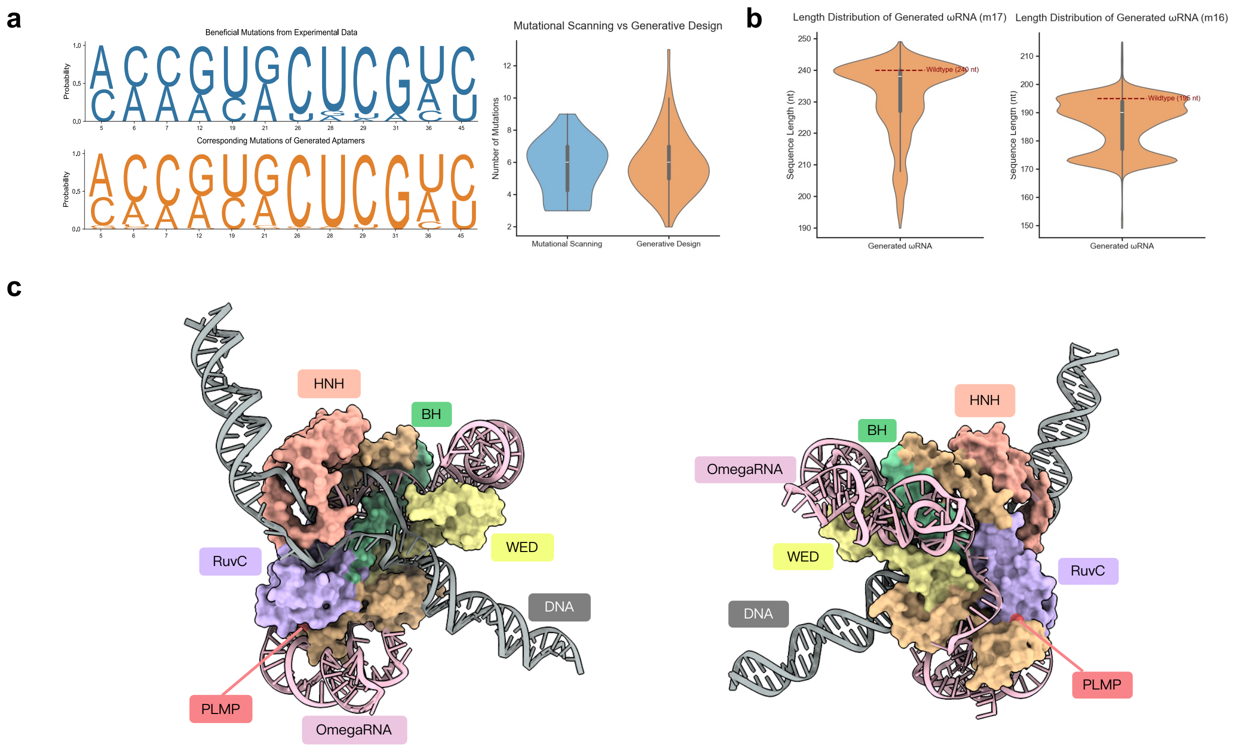


Figure S8: **Fine-tuning enables controlled de novo design of RNA aptamers and IscB omegaRNAs.** (a) *Left:* Position-wise comparison of mutation enrichment between de novo generated aptamer sequences and experimentally beneficial mutations identified by deep mutational scanning (DMS). Each position's mutation logo shows the frequency of nucleotide substitutions in EVA-generated sequences (top) versus those associated with improved fluorescence activity (bottom). The overlap demonstrates that fine-tuning steers the model to preferentially generate the same mutations that experimentally confer functional benefit. *Right:* Distribution of the total number of mutated positions per sequence for EVA-generated aptamers ($n = 93$) versus DMS-characterized variants. The $y$-axis indicates the number of mutated positions relative to wildtype; the $x$-axis has no intrinsic ordering and serves only to separate the two groups. Generated sequences span a comparable range of edit distances as the DMS library, demonstrating that fine-tuned generative models produce *de novo* aptamers with biologically reasonable divergence from wildtype–challenging the notion that generative design is inherently less controllable than point-mutation-based approaches. (b) Length distribution of $\omega$RNA sequences generated by two EVA models independently fine-tuned on the m16 ($n = 42$) and m17 ($n = 44$) sequence datasets, respectively (m16: $n = 1{,}800$ sequences generated; m17: $n = 1{,}595$ sequences generated). Because IscB omegaRNAs are substantially larger than the guide RNAs of modern Cas systems, a key design goal is to generate functional yet more compact guides; the length distributions shown here reflect the range of sequence lengths produced by the fine-tuned model. (c) AlphaFold3-predicted ternary complex structure (MSA mode, RCSB template 7UTN) of a representative designed $\omega$RNA with IscB protein and its cognate DNA substrate. The right panel shows the same complex rotated by 180° to reveal the opposite face.

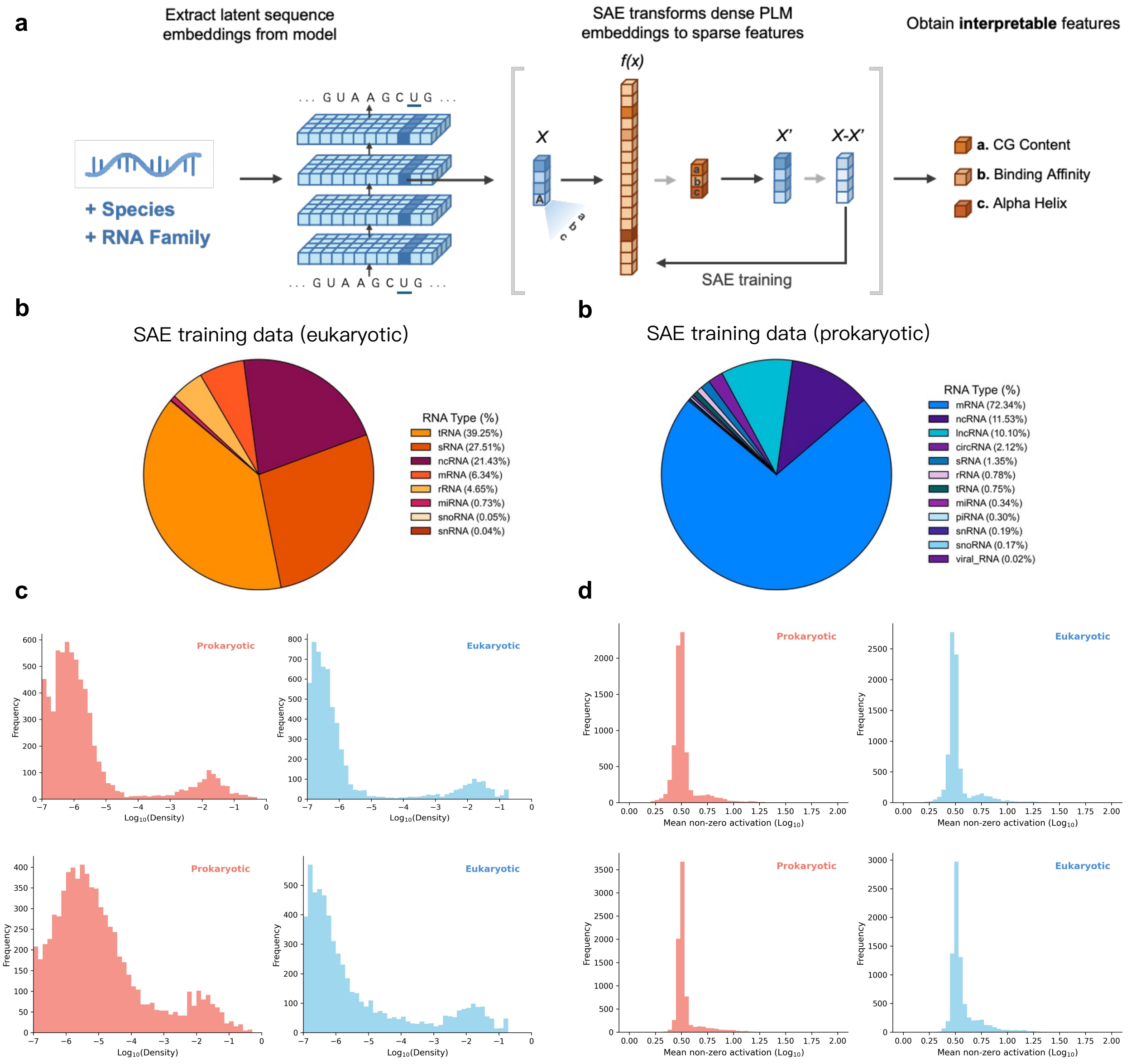


Figure S9: **SAE feature density and activation distributions for prokaryotic and eukaryotic sequences.** (a)(b) SAE feature density distributions ($\log_{10}$) for prokaryotic and eukaryotic sequences without (a) and with (b) lineage prefix prompts. The distributional difference between the two domains becomes more pronounced with the prefix prompt, indicating that lineage conditioning guides the model to invoke distinct SAE feature combinations for prokaryotic and eukaryotic sequences. (c)(d) Mean non-zero activation strength distributions ($\log_{10}$) without (c) and with (d) lineage prefix prompts. Without prompting, eukaryotic sequences already show lower and more uniform activation relative to prokaryotic sequences, which concentrate activation on specific features. This contrast is further amplified by the lineage prefix, demonstrating that species-level context enables more precise discrimination between sequences of different biological origins.

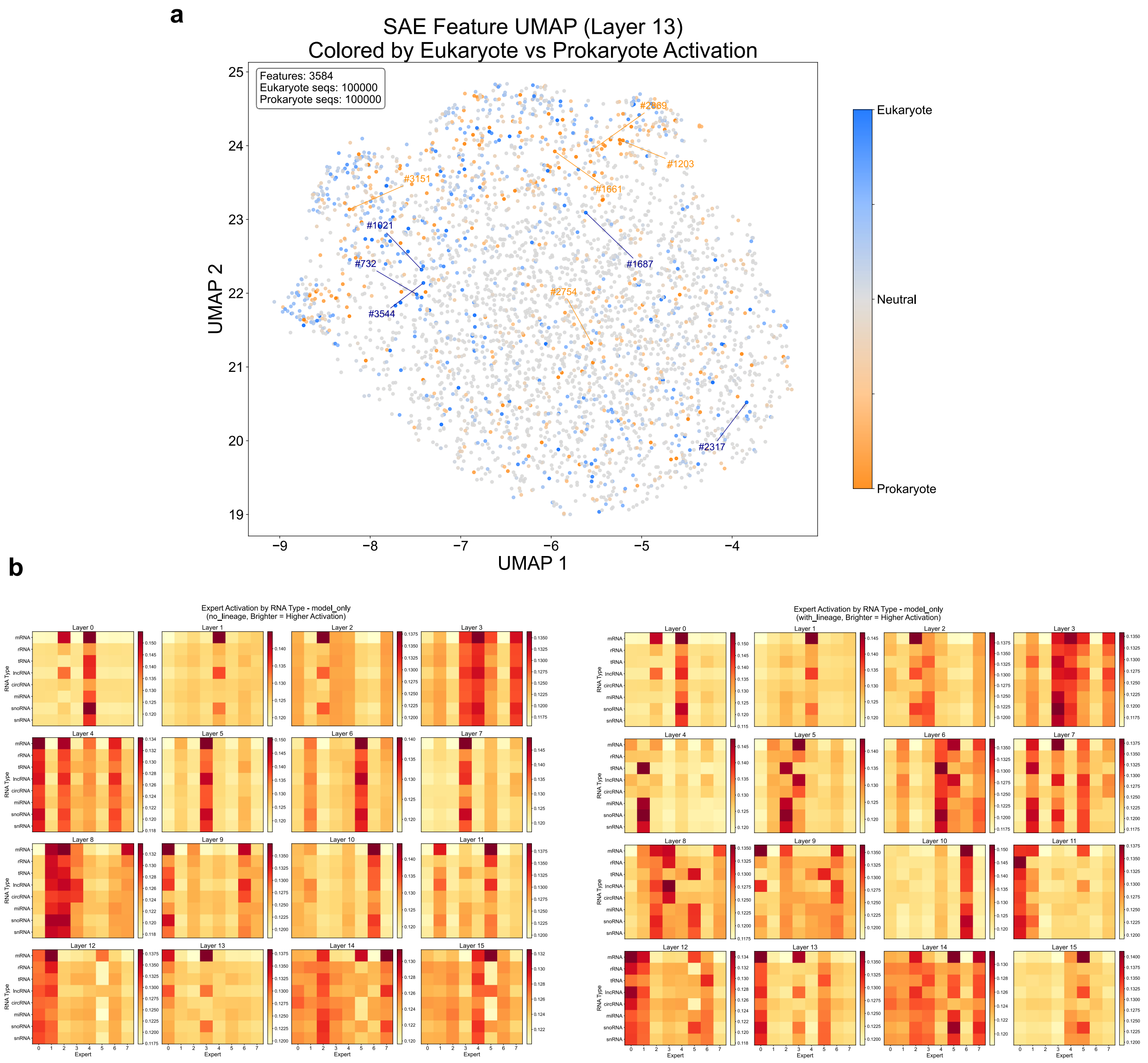


Figure S10: **SAE feature space visualization and MoE expert activation patterns.** (a) UMAP projection of SAE features (3,584 total) colored by differential activation between eukaryotic (blue) and prokaryotic (orange) sequences; gray indicates neutral features. Highly differential features are labeled by index. The clear spatial separation between eukaryote- and prokaryote-preferring features indicates that EVA organizes its internal representations along domain-level biological boundaries. Based on activations from 100,000 sequences per domain. (b)(c) MoE expert activation heatmaps across RNA types (mRNA, lncRNA, rRNA, tRNA, miRNA, circRNA, viral RNA, snRNA, snoRNA) without (b) and with (c) lineage prefix prompts. Brighter colors indicate higher expert activation. Without species context, different RNA types already show subtly distinct expert activation patterns. With lineage conditioning, these differences are substantially sharpened, with RNA types such as rRNA and tRNA developing unique expert activation fingerprints–demonstrating that lineage information enables finer-grained, context-aware routing within the MoE layers.
