## Supplemental tables for "A Long-Context Generative Foundation Model Deciphers RNA Design Principles"

### Supplementary Tables

Table S1: **Comparison of representative RNA foundation models.** “Unified Model”: No if restricted to a narrow RNA subset, Yes if covering mRNA and multiple major ncRNA classes. Context window in nucleotides (nt); “—” = not reported.

| Model | Task | Context(nt) | Unified Model | Training data | Objective |
|---|---|---|---|---|---|
| *Predictive* | | | | | |
| RNA-FM[1] | Predictive | 512 | No (ncRNA) | RNACentral v19; 23M ncRNA | MLM (15%) |
| RiNALMo[2] | Predictive | 1,024 | No (ncRNA) | RNACentral, Rfam, nt, Ensembl; 36M ncRNA | MLM |
| Uni-RNA[3] | Predictive | 1,024 | No (ncRNA) | RNACentral, nt, Genome Warehouse; ∼1B | MLM |
| GARNET[4] | Predictive | 1,024 | No (rRNA) | GTDB genomes (rRNA) | MLM |
| AIDO.RNA[5] | Predictive | 1,024 | No (ncRNA) | RNACentral v24.0; 42M ncRNA | MLM (15%) |
| 5′UTR-LM[6] | Predictive | — | No (mRNA 5′UTR) | 700K 5′UTR seqs | MLM + others |
| HydraRNA[7] | Predictive | 4,096 | Yes (ncRNA + mRNA) | RNACentral, NCBI; 28M ncRNA + mRNA | MLM |
| *Generative* | | | | | |
| SANDSTORM[8] | Generative | <512 | No (toehold, 5′UTR) | — | GAN |
| RNAGEN[9] | Generative | <512 | No (piRNA) | 50,397 natural piRNA | GAN |
| GenerRNA[10] | Generative | 1,024 | No (ncRNA) | RNACentral release 22; 34.39M | CLM |
| RNAGenesis[11] | Generative | 1,024 | Yes (ncRNA+mRNA) | RNACentral; 42M | Diffusion |
| *This work* | | | | | |
| **EVA (ours)** | **Generative** | **8,192** | **Yes (ncRNA + mRNA)** | **OpenRNA v1; 114M full-length RNA** | **CLM + GLM** |

Table S2: **Composition of the OpenRNA v1 training dataset.** Sequence counts before and after quality filtering are reported for each data source, along with the percentage contribution to the final dataset of 114M sequences.

| Data Source | Type | Total Seqs | Filtered Seqs | % | Note |
|---|---|---|---|---|---|
| NCBI (NT & Virus) | Comprehensive | 63,007,114 | 56,232,788 | 49.25% | RefSeq, GenBank, NCBI Virus. |
| RNACentral Consortium | Comprehensive | 51,275,411 | 33,402,433 | 29.25% | RNACentral, Rfam, GtRNAdb, etc. (excl. SILVA). |
| Ensembl | Genomic | 18,739,555 | 15,699,321 | 13.75% | Vertebrate genomes, Release 114. |
| CircRNA Databases | circRNA | 2,219,291 | 2,132,252 | 1.87% | circBase, circAtlas, and others. |
| SILVA | rRNA | 1,312,521 | 552,442 | 0.48% | Ribosomal RNA datasets. |
| NONCODE | lncRNA | 640,747 | 216,709 | 0.19% | Long non-coding RNAs. |
| piRNAdb | piRNA | 200,123 | 41,940 | 0.04% | Piwi-interacting RNAs. |
| Others | Comprehensive | 6,296,930 | 5,908,653 | 5.17% | WormBase, FlyBase, snoDB, miRBase, etc. |
| **Total** | – | **143,691,692** | **114,186,538** | **100%** | Final EVA training set. |

Table S3: **RNA-type sequence conservation and distribution before and after evolutionary conservation-based sampling.** Average Pairwise Identity (API) measures intra-family sequence conservation (Figure S2A); lower values indicate greater sequence diversity. Enrichment ratio = post-sampling proportion / pre-sampling proportion: values $> 1$ indicate up-sampling, values $< 1$ indicate down-sampling (Figure S2D).

| RNA Type | API | Pre-sampling (%) | Post-sampling (%) | Enrichment ratio |
|---|---|---|---|---|
| tRNA | 0.2646 | 5.09 | 1.53 | 0.30 |
| snRNA | 0.2494 | 0.36 | 0.18 | 0.50 |
| rRNA | 0.1361 | 19.89 | 1.08 | 0.05 |
| snoRNA | 0.1041 | 0.39 | 0.17 | 0.44 |
| viral RNA | 0.0991 | 1.49 | 0.25 | 0.17 |
| sRNA | 0.1282 | 5.25 | 1.59 | 0.30 |
| piRNA | 0.1199 | 0.19 | 0.27 | 1.42 |
| miRNA | 0.0759 | 0.78 | 0.36 | 0.46 |
| circRNA | 0.0440 | 1.74 | 1.85 | 1.06 |
| mRNA | 0.0418 | 56.96 | 72.81 | 1.28 |
| lncRNA | 0.0315 | 3.86 | 8.95 | 2.32 |
| ncRNA | – | 4.79 | 10.96 | 2.29 |

Table S4: **Model architecture and training hyperparameters for EVA at different scales.** Architecture, total parameters (Params), active parameters (Active Params), number of layers, model dimension ($d_{\text{model}}$), number of attention heads (Attn Heads), feed-forward network dimension ($d_{\text{FFN}}$), learning rate (LR), weight decay (WD), batch size (BSZ), and warmup steps (WU) are reported for each model variant.

| Arch. | Params | Active Params | Layers | $d_{\text{model}}$ | Heads | $d_{\text{FFN}}$ | LR | WD | BSZ | WU |
|---|---|---|---|---|---|---|---|---|---|---|
| Dense | 32.5M | 32.5M | 16 | 448 | 7 | 1344 | $10^{-4}$ | $5{\times}10^{-6}$ | 16.8M | 3000 |
| Sparse | 21M | 6.5M | 6 | 256 | 8 | 768 | $10^{-4}$ | $5{\times}10^{-6}$ | 1.05M | 3000 |
| Sparse | 31.8M | 11.5M | 11 | 320 | 5 | 480 | $10^{-4}$ | $5{\times}10^{-6}$ | 1.05M | 3000 |
| Sparse | 145M | 51.8M | 16 | 448 | 7 | 1344 | $10^{-4}$ | $5{\times}10^{-6}$ | 2.1M | 3000 |
| Sparse | 437M | 145.1M | 20 | 672 | 12 | 2016 | $10^{-4}$ | $5{\times}10^{-6}$ | 2.1M | 3000 |
| Sparse | 1.4B | 437.2M | 26 | 1024 | 16 | 3072 | $10^{-4}$ | $5{\times}10^{-6}$ | 2.1M | 3000 |

Table S5: **Performance comparison between sparse (MoE) and dense architectures under matched active parameters.** With comparable active parameter counts (∼32M), the sparse MoE model achieves lower validation loss and perplexity (PPL) than the dense model, while requiring fewer FLOPs, demonstrating the efficiency advantage of the MoE architecture for RNA sequence modeling.

| Architecture | Params (Active) | FLOPs | Val Loss | Val PPL |
|---|---|---|---|---|
| Dense | 32.5M | $4.40 \times 10^{20}$ | 1.324 | 3.758 |
| Sparse | 32.4M | $4.36 \times 10^{20}$ | 1.309 | 3.702 |

Table S6: **ncRNA zero-shot fitness benchmark datasets.** All 13 assays were curated from published deep mutational scanning (DMS) studies. Datasets are grouped by RNA class.

| # | Dataset | Source | # Seq |
|---|---|---|---|
| *Ribozyme* | | | |
| 1 | Andreasson_2020_glms | Comprehensive sequence-to-function mapping of cofactor-dependent RNA catalysis in the glmS ribozyme | 3,264 |
| 2 | Janzen_2022_fam1b1 | Emergent properties as by-products of prebiotic evolution of aminoacylation ribozymes | 1,953 |
| 3 | Janzen_2022_fam21 | Emergent properties as by-products of prebiotic evolution of aminoacylation ribozymes | 1,953 |
| 4 | Janzen_2022_fam31 | Emergent properties as by-products of prebiotic evolution of aminoacylation ribozymes | 1,953 |
| 5 | Kobori_2015_ribozyme_j12 | High-throughput assay and engineering of self-cleaving ribozymes by sequencing | 255 |
| 6 | Milena_2021_cata | In vitro selections with RNAs of variable length converge on a robust catalytic core | 135 |
| *RNA Aptamer* | | | |
| 7 | Chen_2024_myo | Bright and stable cyan fluorescent RNA enables multicolor RNA imaging in live *Escherichia coli* | 142 |
| 8 | Zuo_2023_okra | Imaging the dynamics of messenger RNA with a bright and stable green fluorescent RNA | 113 |
| 9 | Chen_2019_pepper | Visualizing RNA dynamics in live cells with bright and stable fluorescent RNAs | 64 |
| 10 | Li_2023_clivia | Large Stokes shift fluorescent RNAs for dual-emission fluorescence and bioluminescence imaging in live cells | 49 |
| *tRNA* | | | |
| 11 | Domingo_2018_tRNA | Pairwise and higher-order genetic interactions during the evolution of a tRNA | 4,175 |
| 12 | Li_2016_tRNA | The fitness landscape of a tRNA gene | 65,536 |
| 13 | Michael_2014_tRNA | Identification of the determinants of tRNA function and susceptibility to rapid tRNA decay by high-throughput in vivo analysis | 213 |
| **Total** | | | **79,805** |

Table S7: **mRNA zero-shot fitness benchmark datasets.** All five datasets were curated from published DMS studies.

| # | Dataset | Source | # Seq |
|---|---|---|---|
| 1 | F7YBW8_MESOW_Ding_2023 | Protein design using structure-based residue preferences | 7,922 |
| 2 | GFP_AEQVI_Sarkisyan_2016 | Local fitness landscape of the green fluorescent protein | 51,714 |
| 3 | Julien_2016_mRNA_rnagym | The complete local genotype-phenotype landscape for the alternative splicing of a human exon | 189 |
| 4 | Ke_2017_mRNA | Saturation mutagenesis reveals manifold determinants of exon definition | 5,560 |
| 5 | Rouskin_2024_mRNA | `https://huggingface.co/datasets/rouskinlab/human_mRNA` | 1,456 |
| **Total** | | | **66,841** |

Table S8: **Human protein DMS datasets used for zero-shot fitness prediction.** Twenty datasets providing readily accessible nucleotide and protein sequences were selected from ProteinGym and DomainOme. For DomainOme entries, the Pfam family identifier and domain position are encoded in the dataset name.

| # | Dataset Name | Source |
|---|---|---|
| 1 | ERBB2_HUMAN_Elazar_2016 | ProteinGym |
| 2 | NPC1_HUMAN_Erwood_2022_RPE1 | ProteinGym |
| 3 | LYAM1_HUMAN_Elazar_2016 | ProteinGym |
| 4 | GLPA_HUMAN_Elazar_2016 | ProteinGym |
| 5 | PITX2_HUMAN_Tsuboyama_2023_2L7M | ProteinGym |
| 6 | RBP1_HUMAN_Tsuboyama_2023_2KWH | ProteinGym |
| 7 | RASH_HUMAN_Bandaru_2017 | ProteinGym |
| 8 | CBPA2_HUMAN_Tsuboyama_2023_1O6X | ProteinGym |
| 9 | PIN1_HUMAN_Tsuboyama_2023_1I6C | ProteinGym |
| 10 | DNJA1_HUMAN_Tsuboyama_2023_2LO1 | ProteinGym |
| 11 | Q9HC78_PF00096_635 | DomainOme |
| 12 | Q9UJQ4_PF00096_383 | DomainOme |
| 13 | Q8N1W1_PF00130_651 | DomainOme |
| 14 | Q9NU63_PF00096_176 | DomainOme |
| 15 | Q7Z5Q1_PF16366_513 | DomainOme |
| 16 | Q9ULZ3_PF00619_114 | DomainOme |
| 17 | Q13263_PF00628_620 | DomainOme |
| 18 | Q9UL15_PF02179_280 | DomainOme |
| 19 | O15151_PF00641_299 | DomainOme |
| 20 | Q9UJQ4_PF00096_627 | DomainOme |

Table S9: **Species used in the zero-shot gene essentiality benchmark.** Both bacterial and eukaryotic essentiality annotations were sourced from the Database of Essential Genes (DEG)[12], cross-referenced with NCBI RefSeq genome annotations. 42 bacterial species were retained if they contained at least 10 essential and 10 non-essential annotated genes. 5 eukaryotic species were assembled from DEG essentiality annotations.

| **Domain** | **Accession** | **Species** |
|---|---|---|
| Bacteria (42) | NC_009085 | *Acinetobacter baumannii* ATCC 17978 |
| | NC_005966 | *Acinetobacter baylyi* ADP1 |
| | NC_000964 | *Bacillus subtilis* subsp. *subtilis* str. 168 |
| | NC_014171 | *Bacillus thuringiensis* BMB171 |
| | NC_016776 | *Bacteroides fragilis* 638R |
| | NC_004663 | *Bacteroides thetaiotaomicron* VPI-5482 |
| | NC_014375 | *Brevundimonas subvibrioides* ATCC 15264 |
| | NC_006350 | *Burkholderia pseudomallei* K96243 |
| | NC_007650 | *Burkholderia thailandensis* E264 |
| | NC_008787 | *Campylobacter jejuni* subsp. *jejuni* 81-176 |
| | NC_002163 | *Campylobacter jejuni* subsp. *jejuni* NCTC 11168 |
| | NC_011916 | *Caulobacter vibrioides* NA1000 |
| | NC_000913 | *Escherichia coli* str. K-12 substr. MG1655 |
| | NC_008601 | *Francisella tularensis* subsp. *novicida* U112 |
| | NC_000907 | *Haemophilus influenzae* Rd KW20 |
| | NC_000915 | *Helicobacter pylori* 26695 |
| | NC_000962 | *Mycobacterium tuberculosis* H37Rv |
| | NC_000908 | *Mycoplasmoides genitalium* G37 |
| | NC_002771 | *Mycoplasmopsis pulmonis* UAB CTIP |
| | NC_010729 | *Porphyromonas gingivalis* ATCC 33277 |
| | NC_008463 | *Pseudomonas aeruginosa* UCBPP-PA14 |
| | NC_009511 | *Rhizorhabdus wittichii* RW1 |
| | NC_005296 | *Rhodopseudomonas palustris* CGA009 |
| | NC_004631 | *Salmonella enterica* serovar Typhi str. Ty2 |
| | NC_016856 | *Salmonella enterica* serovar Typhimurium str. 14028S |
| | NC_003197 | *Salmonella enterica* serovar Typhimurium str. LT2 |
| | NC_016810 | *Salmonella enterica* serovar Typhimurium str. SL1344 |
| | NC_004347 | *Shewanella oneidensis* MR-1 |
| | NC_002952 | *Staphylococcus aureus* MRSA252 |
| | NC_002953 | *Staphylococcus aureus* MSSA476 |
| | NC_003923 | *Staphylococcus aureus* MW2 |
| | NC_002745 | *Staphylococcus aureus* N315 |
| | NC_007795 | *Staphylococcus aureus* NCTC 8325 |
| | NC_010079 | *Staphylococcus aureus* USA300_TCH1516 |
| | NC_007432 | *Streptococcus agalactiae* A909 |
| | NC_003098 | *Streptococcus pneumoniae* R6 |
| | NC_003028 | *Streptococcus pneumoniae* TIGR4 |
| | NC_007297 | *Streptococcus pyogenes* MGAS5005 |
| | NC_011375 | *Streptococcus pyogenes* NZ131 |
| | NC_009009 | *Streptococcus sanguinis* SK36 |
| | NC_007595 | *Synechococcus elongatus* PCC 7942 |
| | NC_002506 | *Vibrio cholerae* O1 biovar El Tor str. N16961 |
| Eukaryota (5) | NC_003070 | *Arabidopsis thaliana* |
| | NC_007194 | *Aspergillus fumigatus* Af293 |
| | NC_003279 | *Caenorhabditis elegans* |
| | NC_001133 | *Saccharomyces cerevisiae* S288C |
| | NC_003421 | *Schizosaccharomyces pombe* |

Table S10: **Definition of repetitive fragments used as a generation quality filter.** Repetitive fragments are identified as unit motifs of length 1–6 bp repeated at least 4 consecutive times. A total of 5,440 detectable combinations are used to screen generated sequences.

| Unit length | Detectable combinations | Min. detection length | Example |
|---|---|---|---|
| 1 bp (Homopolymer) | 4 | 8 bp | AAAAAAAA |
| 2 bp | 12 | 8 bp | AUAUAUAU |
| 3 bp | 60 | 12 bp | AUCAUCAUCAUC |
| 4 bp | 252 | 16 bp | AUCGAUCGAUCGAUCG |
| 5 bp | 1020 | 20 bp | AUUUCAUUUCAUUUCAUUUC |
| 6 bp | 4092 | 24 bp | AUUUUCAUUUUCAUUUUCAUUUUC |
| **Total** | 5440 | — | |

Table S11: **KL divergence KL(Gen‖Nat) comparison between EVA and GenerRNA across 11 RNA classes.** Three secondary-structure feature groups are reported: MFE/GC content (MFE & GC), loop/helix composition (Loop/Helix), and base-pairing rate (Pair/Stem). Lower values indicate better agreement with natural distributions. The Sum column gives the total KL divergence across all three feature groups. EVA (this work) outperforms GenerRNA [10] across all RNA families.

| | GenerRNA | | | | EVA (ours) | | | |
|---|---|---|---|---|---|---|---|---|
| **RNA Type** | **MFE & GC** | **Loop/Helix** | **Pair/Stem** | **Sum** | **MFE & GC** | **Loop/Helix** | **Pair/Stem** | **Sum** |
| mRNA | 1.5725 | 16.9560 | 1.3936 | 19.9221 | 0.4548 | 0.2866 | 0.3538 | **1.0951** |
| circRNA | 1.1211 | 11.3342 | 0.4875 | 12.9429 | 0.4802 | 0.2599 | 0.4362 | **1.1763** |
| lncRNA | 1.4135 | 14.3025 | 0.7218 | 16.4378 | 0.2008 | 1.0691 | 0.1628 | **1.4327** |
| miRNA | 4.2934 | 2.1675 | 5.9571 | 12.4181 | 0.4200 | 0.7216 | 0.7277 | **1.8693** |
| piRNA | 15.6271 | 10.0290 | 15.4934 | 41.1495 | 0.1706 | 0.2007 | 0.2779 | **0.6493** |
| rRNA | 0.5388 | 7.3480 | 0.3635 | 8.2503 | 0.3155 | 0.5712 | 0.2301 | **1.1167** |
| sRNA | 1.1782 | 4.2741 | 0.8189 | 6.2711 | 0.1327 | 0.1754 | 0.1314 | **0.4395** |
| snRNA | 6.7815 | 8.4053 | 5.8128 | 20.9995 | 0.3333 | 0.1618 | 0.2817 | **0.7768** |
| snoRNA | 8.0028 | 6.2184 | 8.3725 | 22.5937 | 0.3998 | 0.1440 | 1.1764 | **1.7202** |
| tRNA | 6.3225 | 5.2746 | 5.4935 | 17.0906 | 0.2468 | 0.2276 | 0.1989 | **0.6733** |
| RNA virus | 2.1901 | 2.1236 | 1.0995 | 5.4132 | 0.2631 | 2.9192 | 0.2422 | **3.4245** |
| **Total** | 48.6015 | 88.0132 | 46.0061 | **182.4988** | 3.5189 | 6.9779 | 4.4220 | **14.2739** |

Table S12: **KL divergence KL(Gen‖Nat) comparison between EVA and GenerRNA for mRNA across six representative species.** Three secondary-structure feature groups are reported: MFE/GC content (MFE & GC), loop/helix composition (Loop/Helix), and base-pairing rate (Pair/Stem). Lower values indicate better agreement with natural distributions. The Sum column gives the total KL divergence across all three feature groups. EVA (this work) with species conditioning outperforms GenerRNA[10] across different organisms.

| | **GenerRNA** | | | | **EVA (ours)** | | | |
|---|---|---|---|---|---|---|---|---|
| **Species** | **MFE & GC** | **Loop/Helix** | **Pair/Stem** | **Sum** | **MFE & GC** | **Loop/Helix** | **Pair/Stem** | **Sum** |
| *C. elegans* | 2.8801 | 18.3659 | 1.1270 | 22.3730 | 0.1073 | 0.1561 | 0.0747 | **0.3380** |
| *C. griseus* | 1.8814 | 18.3367 | 1.4888 | 21.7070 | 0.3964 | 0.2069 | 0.2601 | **0.8634** |
| *H. sapiens* | 1.2351 | 18.3742 | 1.3162 | 20.9254 | 0.2493 | 0.2831 | 0.2255 | **0.7579** |
| *D. melanogaster* | 2.3322 | 18.3063 | 1.5728 | 22.2113 | 0.1124 | 0.0869 | 0.0739 | **0.2732** |
| *M. musculus* | 1.3512 | 18.3707 | 1.1570 | 20.8789 | 0.4706 | 0.3367 | 0.2713 | **1.0785** |
| *R. norvegicus* | 1.6329 | 18.3360 | 1.2585 | 21.2274 | 0.3232 | 0.5825 | 0.1531 | **1.0588** |
| **Total** | 11.3129 | 110.0898 | 7.9204 | **129.3230** | 1.6592 | 1.6522 | 1.0586 | **4.3699** |

Table S13: **Domain-level structural criteria for evaluating IscB–ωRNA complex plausibility.** Criteria were derived from published structural analyses of the IscB–ωRNA–DNA ternary complex[13,14].

| Domain | Biological Function | Structural Criterion | Metric / Threshold |
|---|---|---|---|
| HNH | Nuclease domain responsible for cleaving the target DNA strand (the strand complementary to the ωRNA spacer). | Distance between the HNH domain catalytic residues and the guide RNA is measured to confirm productive positioning of the nuclease relative to the RNA–DNA hybrid. | HNH–ωRNA Cα–P distance |
| PLMP | A four-residue motif that contacts both the RuvC domain and the ωRNA, stabilizing the overall IscB–ωRNA complex architecture. | All four PLMP residues must make direct contact with the ωRNA; a residue is considered in contact if any heavy-atom distance to the RNA falls below the threshold. | All 4 residues $< 7$ Å from ωRNA |
| BH straightness | The bridge helix is an α-helical element that traverses the full length of the ωRNA channel, forming the central scaffold of the complex. | Helical straightness of the BH is assessed by the deviation of Cα atoms from the principal helix axis; excessive bending indicates scaffold disruption. | BH Cα straightness index |
| BH wrap-around | The guide RNA encircles the bridge helix, and sufficient wrap-around coverage is required for stable complex assembly. | Fraction of the BH surface covered by the encircling ωRNA; low coverage indicates loss of the central guide-RNA scaffold. | ωRNA wrap-around coverage fraction |
| WED | Mediates orthogonal recognition of the ωRNA scaffold and interacts with the phosphate backbone of the PAM-proximal region of the target DNA. | Distance from the WED domain to the PAM-proximal DNA backbone is measured to verify appropriate positioning for PAM-dependent target engagement. | WED–PAM backbone distance |

Table S14: **Shared UTR sequences used in all four mRNA vaccine constructs.**

| Region | Length | Sequence |
|---|---|---|
| 5′UTR | 64 nt | AGGAAAUUCCAUUUGGCUGCAGCUUCUGGAGGGAGCCGACAGGAGACGUGGGGAGACGGCCACC |
| 3′UTR | 94 nt | GCUGCCUUCUGCGGGGCUUGCCUUCUGGCCAUGCCCUUCUUCUCUCCCUU<br>GCACCUGUACCUCUUGGUCUUUGAAUAAAGCCUGAGUAGGAAGU |

Table S15: **mRNA vaccine codon optimization results.** Results are sorted by CAI in descending order within each vaccine system. MFE: minimum free energy; CAI: codon adaptation index; Acceptance Rate: fraction of synonymous substitutions accepted during optimization. Protein sequences, original and optimized RNA sequences are provided in the Supplementary Data.

| **Vaccine** | **Model** | **CAI** | **MFE (kcal/mol)** | **Accept. Rate** |
|---|---|---|---|---|
| **HIV-1 gp160** | EVA (lineage: Human) | 0.8458 | −1168.50 | 55.6% |
| GenBank K03455.1 | EVA (no lineage) | 0.7931 | −1101.90 | 62.2% |
| CDS: 2571 nt (856 aa) | CodonFM-1B | 0.7908 | −1059.00 | 51.9% |
| mRNA: 2729 nt | Evo2-1B | 0.7564 | −1099.60 | 45.6% |
| **Influenza PR8 HA** | EVA (no lineage) | 0.9115 | −904.80 | 73.3% |
| A/PR/8/34 (H1N1) | EVA (lineage: Human) | 0.9087 | −894.40 | 61.3% |
| CDS: 1701 nt (566 aa) | CodonFM-1B | 0.8877 | −895.70 | 66.2% |
| mRNA: 1859 nt | Evo2-1B | 0.7711 | −756.90 | 50.0% |
| **Rabies RABV-G** | EVA (lineage: Human) | 0.9056 | −910.80 | 56.9% |
| GenBank GQ918139 | EVA (no lineage) | 0.8756 | −924.60 | 71.3% |
| CDS: 1575 nt (524 aa) | CodonFM-1B | 0.8310 | −935.30 | 55.6% |
| mRNA: 1733 nt | Evo2-1B | 0.7782 | −889.30 | 58.1% |
| **VZV gE** | EVA (lineage: Human) | 0.8692 | −1034.90 | 64.4% |
| Varicella-Zoster Virus | EVA (no lineage) | 0.7878 | −1031.40 | 64.4% |
| CDS: 1749 nt (582 aa) | Evo2-1B | 0.7404 | −1012.10 | 50.0% |
| mRNA: 1907 nt | CodonFM-1B | 0.7402 | −1067.80 | 51.9% |
| | EVA (lineage: Human) | 0.8422 | −585.42 | 60.6% |
| **SARS-CoV-2 Spike** | EVA (no lineage) | 0.8150 | −603.40 | 66.9% |
| Wuhan-Hu-1 (NC_045512) | CodonFM-1B | 0.8070 | −590.20 | 53.8% |
| mRNA:1055nt | Evo2-1B | 0.7848 | −516.72 | 38.1% |
| | Random | 0.7411 | −556.70 | 10.0% |

*Note: Protein sequences, original RNA sequences, and optimized RNA sequences for each vaccine are provided in the Supplementary Data.*

Table S16: **circRNA vaccine ORF codon optimization results.** Results are sorted by CAI in descending order within each vaccine system. MFE: minimum free energy (full circRNA construct, computed by ViennaRNA); CAI: codon adaptation index; Acceptance Rate: fraction of synonymous substitutions accepted during optimization. Protein sequences, original and optimized ORF sequences are provided in the Supplementary Data.

| Vaccine | Model | CAI | MFE (kcal/mol) | Accept. Rate |
|---|---|---|---|---|
| **Circ_CAR-T** CircRNA vaccine antigen CDS: 1500 nt (500 aa) | EVA (lineage: Human) | 0.8961 | −1398.80 | 66.9% |
| | EVA (no lineage) | 0.8820 | −1423.30 | 63.1% |
| | CodonFM-1B | 0.8637 | −1425.40 | 65.0% |
| | Evo2-1B | 0.8490 | −1356.70 | 59.4% |
| | Random | 0.7579 | −1315.90 | 16.9% |
| **Circ_RABV** Rabies Virus Glycoprotein G CDS: 1574 nt (524 aa) | EVA (lineage: Human) | 0.8589 | −1069.10 | 65.0% |
| | EVA (no lineage) | 0.8420 | −1077.60 | 52.5% |
| | CodonFM-1B | 0.8298 | −1121.40 | 60.0% |
| | Evo2-1B | 0.8036 | −1042.20 | 62.5% |
| | Random | 0.7176 | −1007.60 | 18.1% |
| **Circ_SARS (Group-I)** SARS-CoV-2 Spike Group-I intron cyclization CDS: 669 nt (223 aa) | EVA (lineage: Human) | 0.8592 | −870.60 | 59.4% |
| | EVA (no lineage) | 0.8578 | −869.70 | 65.0% |
| | Evo2-1B | 0.7644 | −823.60 | 34.4% |
| | CodonFM-1B | 0.7441 | −871.70 | 76.2% |
| | Random | 0.7314 | −846.90 | 11.9% |
| **Circ_SARS (T4 ligase)** SARS-CoV-2 Spike T4 RNA ligase cyclization CDS: 669 nt (223 aa) | EVA (lineage: Human) | 0.8822 | −719.30 | 56.2% |
| | EVA (no lineage) | 0.8344 | −715.40 | 58.8% |
| | Evo2-1B | 0.7717 | −675.90 | 35.6% |
| | CodonFM-1B | 0.7567 | −722.40 | 66.2% |
| | Random | 0.6911 | −708.00 | 9.4% |

*Note: Protein sequences, original ORF sequences, and optimized ORF sequences for each circRNA vaccine are provided in the Supplementary Data.*
